# Neuro-Metabolic and Vascular Dysfunction as an Early Diagnostic for Alzheimer’s Disease and Related Dementias

**DOI:** 10.1101/2025.05.12.653558

**Authors:** Juan Antonio K. Chong Chie, Scott A. Persohn, Ravi S. Pandey, Olivia R. Simcox, Paul Salama, Paul R. Territo, the Alzheimer’s Disease Neuroimaging Initiative

## Abstract

Alzheimer’s disease (AD) is the most prevalent neurodegenerative condition characterized by significant cognitive decline. Recent studies suggest that the brain undergoes anatomical and functional restructuring, resulting in neuro-metabolic and vascular dysregulation (MVD) prior to amyloid-β accumulation, which begins at an early age and leads to the onset of AD. Using a retrospective clinical population (N=403) of subjects with varying disease stages from the Alzheimer’s Disease Neuroimaging Initiative (ADNI), we identified that disease progression follows a stage-dependent MVD pattern, facilitating the identification of at-risk and resilient brain regions. Although each region progresses at a different pace, regions associated with memory, cognitive tasks, and motor function showed significant early dysregulation. These changes aligned with transcriptomics and cognitive signatures. This study underscores that MVD in brain regions varies by sex and disease stage, making it a sensitive tool for early AD diagnosis. Furthermore, this approach could improve patient monitoring, stratification, and therapeutic testing.

---

Alzheimer’s disease (AD) and related dementia (RD) is the fifth-leading cause of death in the USA and the most common cause of dementia, with an economic burden of more than a trillion US dollars annually.^1^ ADRD is characterized by gradual neurodegeneration, which leads to a decline in cognitive function.^1^ Clinically, patients and/or caregivers seek neurological assessments; however, by this point neurodegeneration has reached an irreversible state.^1^ This temporal gap between symptom onset and clinical diagnosis highlights the need for early and accurate diagnosis of neurodegenerative disorders. Efforts to improve diagnostic methods using medical imaging, such as positron emission tomography (PET) and magnetic resonance imaging (MRI), aim to predict disease onset by detecting early functional and structural changes before significant amyloid-β accumulation, which begins 20 to 30 years prior to symptoms.^1,2^ Historically, AD diagnosis via imaging has focused on the evaluation of late-stage, single modality, and structural or functional changes, which lack sensitivity and specificity during neurodegeneration.^2,3^ Consequently, treatment planning based on these low fidelity readouts results in outcomes that are compromised, leading to ineffective therapeutic responses.^2,3^

The brain is the most metabolically active organ in the body, consuming about 20% of the body’s total energy at rest (i.e. 260–320 calories) per day.^4^ Due to the limited metabolic buffering capacity of the brain, this energy is supplied via approximately 15-20% of the total cardiac output of the heart under resting conditions.^4^ Based on this, the neuro-metabolic requirements must be met via coordinated increases in regional blood flow to prevent energy deficits,^4^ and is referred to as neurovascular coupling (NVC).^4^ Under pathophysiological states, NVC can become dysregulated,^4^ thus leading to disease onset and progression.^4^ Until recently, the assessment of NVC as an early and sensitive diagnostic of ADRD has largely been overlooked.^5^

Several lines of evidence suggest that neurovascular dysregulation begins at an early age, and leads to onset of AD.^5,6^ Moreover, the metabolic reprogramming theory, a bioenergetic model of metabolic regulation between neurons and astrocytes during aging, suggests that metabolic dysregulation and bioenergetics deficits are critical elements in the etiology of the disease.^7^ Additionally, recent preclinical studies from our lab suggest that regional metabolic and vascular dysfunction (MVD) persist across the disease spectrum;^8^ however, few studies have explored these combined measures clinically in AD.^9^ Hence, measuring the interdependence of NVC and its dysregulation in an individual, when compared to population norms, could provide valuable insights into disease onset, and how this progresses across the AD spectrum.

To this end, recent evidence suggests that cerebral metabolic and perfusion alterations are associated with a gliosis in response to metabolic insufficiency and oxidative stress,^7^ and driven by chronic inflammation.^10^ Importantly, this leads to a feed forward regulation of this cascade, resulting in transcriptomic alteration in metabolic and vascular gene ontology signatures,^11^ microglial activation,^12^ increased neuronal oxidative stress,^13^ astrocytosis,^14^ and enhanced MVD.^8,9^ To maintain neuro-metabolic function during these periods, it has been suggested the neuronal glycolytic insufficiency is supplemented by astroglia metabolism and shuttling of the monocarboxylates, such as lactate, pyruvate, and ketones via the monocarboxylate transporters (MCTs).^4,14^ Based on this, our lab and others, theorized that astrocyte signaling occurs prior to cognitive impairments become evident.^14^

Therefore, in this work, we hypothesize that AD progression follows a prescribed sequence of MVD changes that are initiated by neuroinflammation, which results in an increase in cerebral blood flow (CBF) as a compensatory response to neuro-metabolic insufficiency, termed Type 1 Uncoupling (T1U). We further hypothesize that sustained inflammation coupled with energy deficits, leads to micro- and astro-gliosis to support MCT mediated energy shuttle to neurons; however, this occurs in excess of CBF, leading to a supply/demand imbalance, termed Type 2 Uncoupling (T2U). Additionally, we hypothesize that this net energy imbalance leads to an increase in CBF (hyper-perfusion) to meet the brain’s heightened energy demands (hyper-metabolism) that are driven in response to rising oxidative stress, a phase termed Prodromal Coupling (PC). Lastly, we hypothesize that during the prodromal phase, the brain reaches a bioenergetic equilibrium, where metabolism and perfusion reach a quasi-stable metabolic state; however, this neuro-metabolic demand exceeds the brains functional reserve capacity, leading to neuronal stress and cell death observed in late AD, resulting in hypo-metabolic and hypo-perfused phase, termed Neuro-Metabolic and Vascular Failure (NMVF). Importantly, we believe this MVD sequence of events provides a deeper understanding of brain activity during AD progression (Figure 1.A).

**Figure 1.**
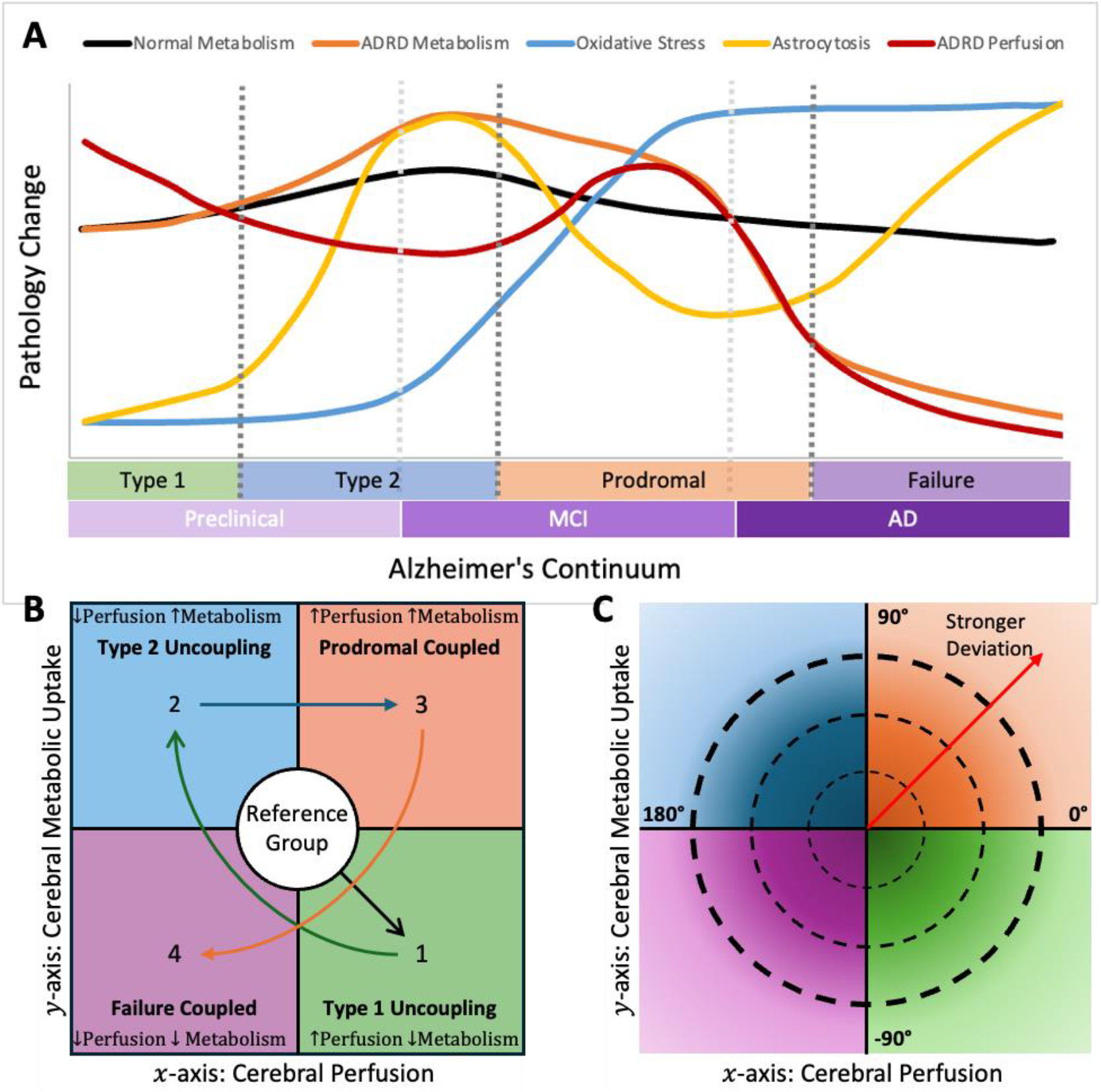
Neuro-vascular and Metabolic Changes Across the AD Spectrum, MVD Pattern, and Polar Coordinates Representation. The normal metabolic state continuum in healthy individuals is depicted by the black line. In contrast, the metabolic state across AD is represented by the orange line, illustrating the metabolic alterations observed throughout the disease’s various phases, as proposed by the metabolic reprogramming theory.^7^ The red line represents the progression of perfusion across the disease, demonstrating how metabolism and perfusion become dysregulated until they eventually fail. During the hypermetabolic phase of the disease, oxidative phosphorylation undergoes upregulation as a compensatory response, leading to progressively oxidative damage. Furthermore, it is suggested that astrocytosis signaling precedes the onset of cognitive decline.^5,6^ (**B**) Neurovascular and metabolic dysregulation were assessed by projecting the z-score SUVR for each brain region onto Cartesian space. The x-axis represents the z-score changes in cerebral perfusion derived from CBF maps, while the y-axis represents the z-score change in glucose uptake derived from FDG. Brain regions toward the center of the plane are likely functioning at a normal level. A brain region is considered coupled if both signals are in the same direction (Cartesian quadrants 1 and 3) and uncoupled if they are opposite (Cartesian quadrants 2 and 4). We hypothesize the existence of a regional disease pathway (MVD pattern) that maps the disease progression and can accurately discern the MVD changes, which is: Type 1 Uncoupling *→* Type 2 Uncoupling *→* Prodromal *→* Neuro-metabolic-vascular Failure. (**C**) Alternatively, Cartesian space readouts can be transformed into Polar coordinates to visualize disease progression while preserving the MVD state. The distance from the origin represents the magnitude of the change, while the angle indicates the deviation of the change.

In this study, we tested the aforementioned hypotheses in a retrospective clinical population containing cognitively normal (CN), early mild cognitive impairment (EMCI), mild cognitive impairment (MCI), late MCI (LMCI), and AD subjects, and demonstrated that MVD analysis serves as a sensitive and early noninvasive diagnostic tool, which can discriminate between stages of AD progression.

## 1. Results

The study population (N=403) was on average 73.56±7.40 years old. Most of the study population was composed of APOE3 homozygotes(ε3/ε3;48.64%) and overweight(46.65%) individuals (For detailed demographic stats, see Table S3). Registration quality assurance was performed for all images by a team of experts to ensure proper alignment and scan coverage. Image alignment quality was measured via structural similarity index measure^15^ across all modalities, which was 0.9143±0.0288 (n=1612,N=403).

### 1.1 Metabolic and Perfusion Analysis

Our lab has developed a framework to assess MVD,^8^ by projecting CBF and FDG z-score readouts onto a Cartesian space, where regional measures fall into one of four different phenotypes (Figure 1.B):

- Type 1 uncoupling (T1U), in Cartesian quadrant 4 (↓FDG,↑CBF), shows a pathophysiologic phenotype with an increase in perfusion and decrease in metabolism. This decrease in metabolism is driven by neuroinflammatory mediators, while the increase in perfusion is driven by reactive hyperemia to offset the energy deficit.^10,16,17^
- Type 2 uncoupling (T2U), in Cartesian quadrant 2 (↑FDG,↓CBF), shows a pathophysiologic phenotype with a decrease in perfusion and increasing metabolism. In this phenotype, chronic neuroinflammation results in vascular dysregulation, while brain metabolism increases are driven by activation of glial cells (i.e. microglia and astrocytes).^12,17^
- Prodromal coupling (PC), in Cartesian quadrant 1 (↑FDG,↑CBF), shows a phenotype where both perfusion and metabolism are increasing, driven by microgliosis in response to accumulation of amyloid plaques and tau tangles,^18,19^ astrogliosis in response to increasing neuronal metabolic demand,^14^ and vascular proliferation driven by angiogenic mediators.^20^
- Neuro-metabolic-vascular failure (NMVF), in Cartesian quadrant 3 (↓FDG,↓CBF), shows both perfusion and metabolism are decreasing due to regional insufficiency resulting in sustained damage.^7,16^

Using this framework, we assessed regional metabolism and perfusion changes in a retrospective population to determine stage-dependent changes in EMCI, MCI, LMCI, and AD subjects relative to the CN. Consistent with our overarching hypothesis, each disease stage showed different magnitudes in perfusion and metabolism alterations, which were sexually dimorphic (Figure 2). For EMCI (Figure 2.A), brain regions of both sexes showed alterations in perfusion and metabolism within one standard deviation relative to the CN group. Male subjects showed significant changes in 21 out of 59 regions (Figure 2.E), 20 of which were either T1U(13) or T2U(7) uncoupled. In contrast, female subjects showed significant changes in 15 out of 59 regions (Figure 2.A), 14 of which were T1U(2) or T2U(12) uncoupled. Unlike EMCI, the MCI group (Figure 2.B) for both sexes showed a more extensive spread of brain regions across the perfusion axis, concomitant with an increase in metabolism, resulting in a larger number of regions in quadrant 2, which retained T2U uncoupling. The number of significant regions in MCI (Figure 2.F) was greater than in EMCI for both sexes (M=29/59, F=30/59), with a large increase in the number of significant regions in T2U (M=15/59, F=13/59). Consistent with our hypothesis, brain regions for both sexes of the LMCI group (Figure 2.C), showed a shift from uncoupled to coupled, where these regions retained either PC or NMVF phenotypes. For males, the number of significant regions (Figure 2.G) was reduced from 29 to 14, when compared to the MCI group. Similarly, females exhibited decrease in the number of significant regions from 30 to 6, compared to the same group. Lastly, for the AD group (Figure 2.D), brain regions for both sexes showed a predominant NMVF phenotype. Both males and females showed an increase in the number of regions (Figure 2.H) that reflect a significant change relative to CN (M=26/59, F=19/59), and a total of 10 brain regions (in both cases) were in the NMVF quadrant.

**Figure 2.**
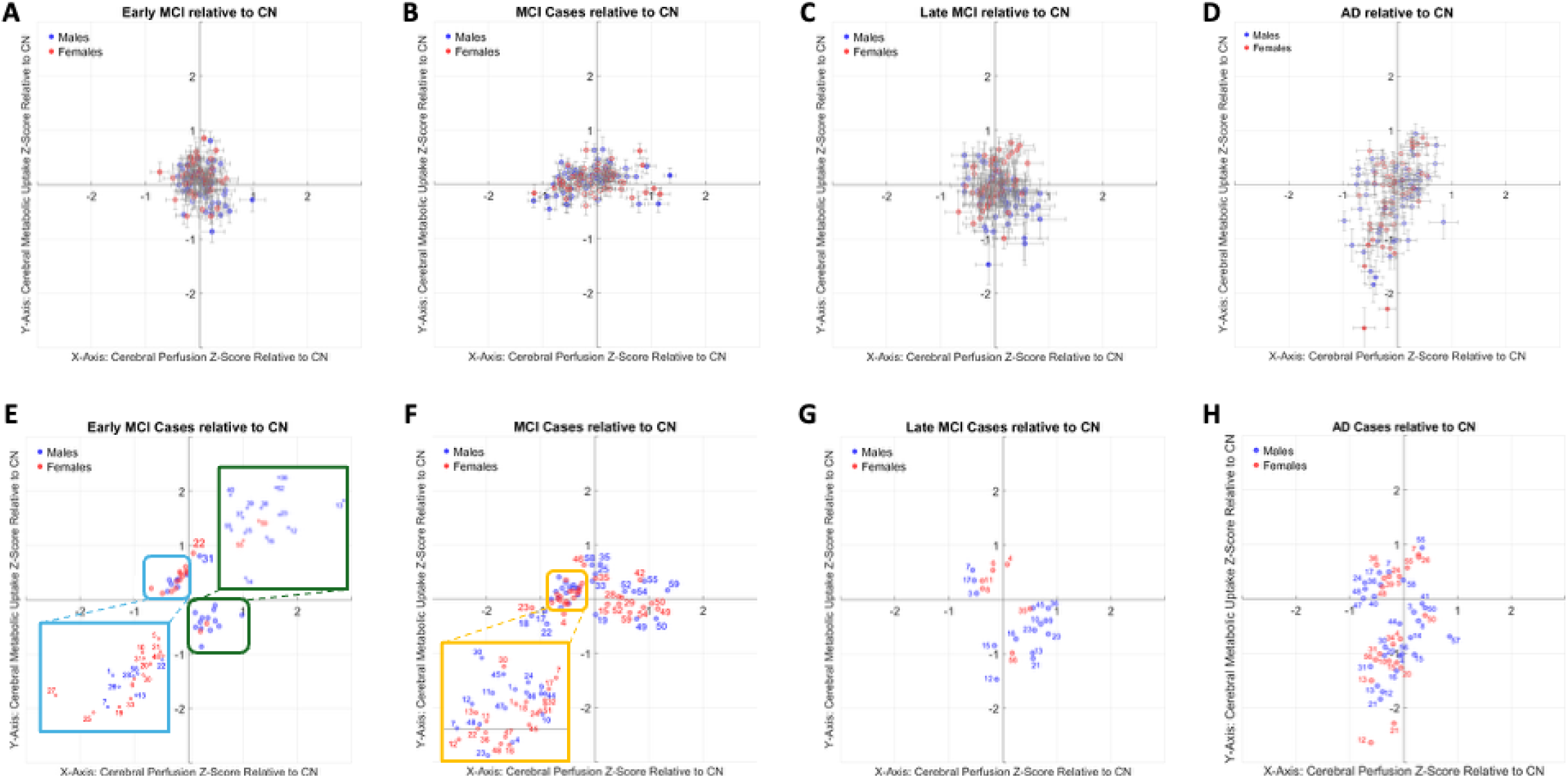
Assessment of Neurovascular Coupling and Uncoupling of EMCI, MCI, LMCI, and AD Relative to CN. Using the MVD framework, each disease stage exhibited distinct alterations in perfusion and metabolism, which were also sexually dimorphic. (**A**) For EMCI, brain regions in male and female subjects showed alterations within one standard deviation of the CN group in both perfusion and metabolism. (**B**) For MCI, both sexes exhibited a more extensive spread of brain regions across the perfusion axis, accompanied by an increase in metabolism. This resulted in a larger number of regions in quadrant 2, which retained T2U uncoupling. (**C**) For LMCI, brain regions in both sexes showed a shift from uncoupled to coupled metabolism. These regions retained either PC or NMVF phenotypes. (**D**) For AD, brain regions in both sexes predominantly exhibited the NMVF phenotype. Significant changes were determined using Student’s t-test (p<0.05). (**E**) For EMCI, males showed significant changes in 21 out of 59 regions and 20 of these regions were either T1U (13) or T2U (7) uncoupled. In contrast, females showed significant changes in 15 out of 59 regions, where 14 of these regions were T1U (2) or T2U (12) uncoupled. (**F**) For MCI, the number of significant regions was greater than the number of significant regions observed for EMCI for both sexes (M=29/59, F=30/59), with a large increase in the number of significant regions in T2U (M=15, F=13). (**G**) For LMCI, the number of significant regions was reduced from 29 to 14 in males, and from 30 to 6 for females, compared to MCI. (**H**) For AD, both males and females showed an increase in the number of regions that reflect a significant change relative to CN (M=26/59, F=19/59), and a total of 10 brain regions (in both cases) were in the NMVF quadrant. Brain regions names and indices can be found in Table S4.

To understand which regions remain significant across disease stages, we followed regional progression in both sexes. Among the 59 unilateral regions, the cingulate gyrus, inferior temporal gyrus, lateral occipital cortex, mid-frontal gyrus, mid-temporal gyrus, occipital pole, precentral gyrus, precuneus cortex, all showed consistently significant disease progression (Figures 2.E-H). By contrast, the cerebellum, frontal pole, frontal medial cortex, pallidum, and superior parietal lobe significantly change only during the first stages of the disease (EMCI/MCI) (Figures 2.E-H). It is worth noting that the hippocampus, frontal operculum cortex, and superior frontal gyrus do not change significantly until late in the disease (LMCI/AD). To illustrate these changes, significant regions were projected into the MNI152+ for spatial localization across all disease stages, by thresholding regions at the p<0.05 level (Figure S1).

### 1.2 Inter-Stage Neuro-Vascular-Metabolic Uncoupling Analysis

To better understand the regional sequence of events in MVD across ADRD stages, we extended our analysis by comparing each stage with its preceding stage to determine the inter-stage changes (e.g. EMCI(CN), MCI(EMCI), LMCI(MCI), AD(LMCI)). During early stages of the disease, EMCI relative to CN (Figure S2.A), showed 21 and 15 significant brain regions for male and females, respectively, which were dominated by T1U. When comparing MCI subjects with EMCI (Figure S2.B), both male and female brain regions demonstrated increased dispersion for both perfusion and metabolism. Since individual brain regions exhibit unique responses, this data suggests that regional demands may drive distinct pathophysiology by neuroinflammation, oxidative stress, and astrocyte activation to maintain baseline function. In the transition from MCI to LMCI (Figure S2.C), we observe a pronounced increase in CBF. However, there was no significant alteration in metabolism over this same interval. These findings support the hypothesis that during this phase, the brain enters a quasi-stable metabolic state by increasing the cerebral blood flow through NVC mechanisms to support the heightened metabolic demand resulting from the energetic imbalance. Finally, when comparing AD subjects with LMCI (Figure S2.D), several brain regions displayed characteristics consistent with the NMVF phenotype, consistent with neurodegenerative process triggered by the prolonged elevated energetic demands of the quasi-stable metabolic phase in excess of the functional capacity. These observations suggest that the brain undergoes sequential vascular and perfusion changes in an effort to balance bioenergetic demands across different disease stages, consistent with our hypotheses.

### 2.3 Uncoupling Migration Charts

To track disease progression across stage, MVD type, and region, we transformed Cartesian plots into polar coordinates space to facilitate visualization. Uncoupling Migration Charts (UMC) permit direct assessment of disease events in the brain from the MVD perspective. The intervals [- 90°,0°),[90°,180°),[0°,90°) and [-180°,-90°) map to the T1U, T2U, PC, and NMVF phenotypes, respectively. The circle’s diameter represents the changes in magnitude, where a larger circle indicates a greater deviation from the reference group. UMC plots for the key brain regions across all disease stages relative to CN are shown in Figure 3 (For the complete plots, see Figures S3-S8). Regions associated with the Mid and Inferior Temporal Gyrus begin as T1U for EMCI subjects, and progress to T2U for MCI; however, by LMCI, these regions have already reached the NMVF phenotype, while in AD, the magnitude of the change increases. Notably, female subjects reach the NMVF phenotype faster for these regions, where the Occipital Pole and the Planum Polare follow a similar trajectory: T1U in EMCI, T2U in MCI, PC in LMCI, and NMVF in AD. Importantly, this progression occurs at different angles and magnitudes, suggesting that each brain regional trajectory is unique. In contrast, certain regions (e.g., Insular Cortex, Frontal Orbital Cortex, Inferior Frontal Gyrus) exhibit minimal to no alterations in magnitude or angle across the disease spectrum.

**Figure 3.**
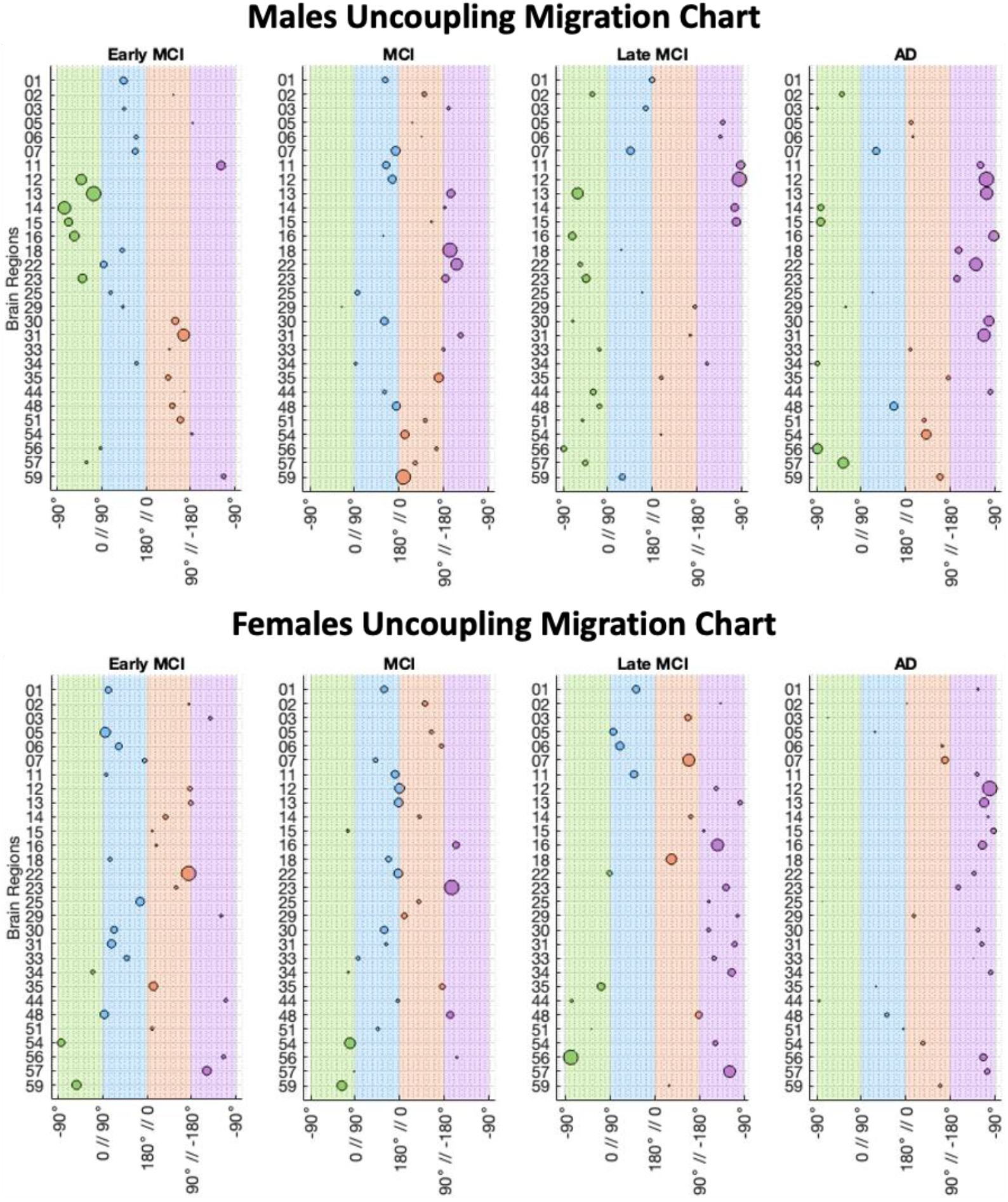
Uncoupling Migration Chart of the Important Brain Regions for each Disease Stages Relative to CN. The uncoupling migration charts show how the brain regions traverse between the uncoupled and coupled phases across disease progression. Changes were measured relative to CN. The intervals [-90°,0°),[90°,180°),[0°,90°) and [-180°,-90°) map directly to the T1U, T2U, PC, and NMVF phenotypes, respectively. The circle’s diameter represents the change’s magnitude, where a larger circle indicates a greater deviation from the reference group. Notably, females reach the NMVF phenotype faster than males; however, this progression occurs at different angles and magnitudes, suggesting that each brain regional trajectory is unique. Brain regions names and indices can be found in Table S4.

### 1.4 Inter-Stage Uncoupling Migration Charts

To understand the sequential progression by stage for these regions, UMC data derived from each disease stage relative to the previous stage are presented in Figure 4 (For the complete plots, see Figures S5 and S8). Compared to the previous stage, regional MVD phenotypes across the disease spectrum were unique, with each region progressing at different rates and magnitudes. For instance, the temporal gyrus regions still follow the trajectory described when compared to the CN population; however, the effect is more pronounced. Additionally, females consistently experience the sequential phases at faster rates, which can be seen in the magnitude of the changes. Notably, the aforementioned regions, which are associated with cognition, memory, and recognition, still reach an NMVF phenotype during AD. However, the regions that exhibit minimal change (such as the Insular Cortex, Frontal Orbital Cortex, and Inferior Frontal Gyrus) still show minimal changes for the analysis relative to the previous stage is performed.

**Figure 4.**
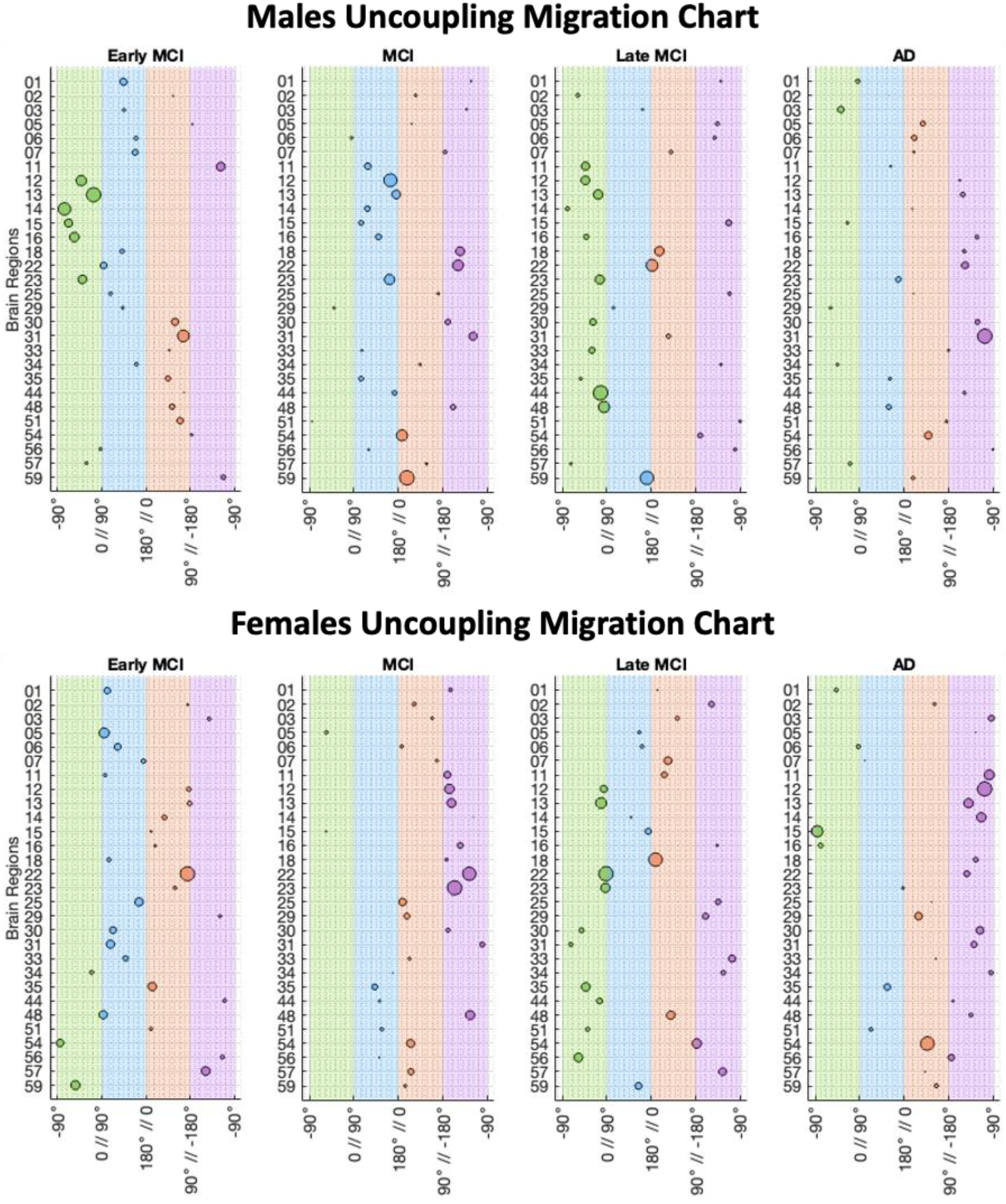
Uncoupling Migration Chart of the Important Brain Regions for each Disease Stages Relative to the Previous Disease Stage. For this uncoupling migration chart, changes were measured relative to the previous stage to understand the sequential inter-stage changes. The intervals [-90°,0°),[90°,180°),[0°,90°) and [-180°,-90°) map directly to the T1U, T2U, PC, and NMVF phenotypes, respectively. The circle’s diameter represents the change’s magnitude, where a larger circle indicates a greater deviation from the reference group. Comparing these plots against the plots relative to CN, significant regions exhibit different MVD phenotypes across the disease spectrum. Moreover, this comparison highlights the varying rates and degrees of effect between each region. Similarly to CN, for AD, the significant regions reach the NMVF phenotype in both sexes. Brain regions names and indices can be found in Table S4.

### 1.5 Transcriptomic Analysis

Analysis of WGCNA^21^ from blood transcriptome identified gene expression changes in a system-level framework, and identified 28 distinct modules of co-expressed genes (Table S5). To elucidate the functional significance of these modules, we correlated each module eigengene with the average z-scores of CBF and FDG imaging data, as well as sex and disease stage (MCI/AD) of the subjects (Figure S9).

We identified two modules significantly associated with CBF (p<0.01; red, grey60) and one module significantly associated with FDG (p<0.05; brown4). We also observed that CBF associated red module and FDG associated brown4 module were significantly correlated (p<0.05) with AD disease stage (Figure 5.A). KEGG pathway enrichment analysis identified enrichment of “porphyrin metabolism” pathway in the red module, multiple disease associated pathways in the skyblue3 module, and “beta-alanine metabolism”, “osteoclast differentiation”, and “toll-receptor signaling pathway” in the brown4 module (Figure 5.B/Table S6).

**Figure 5.**
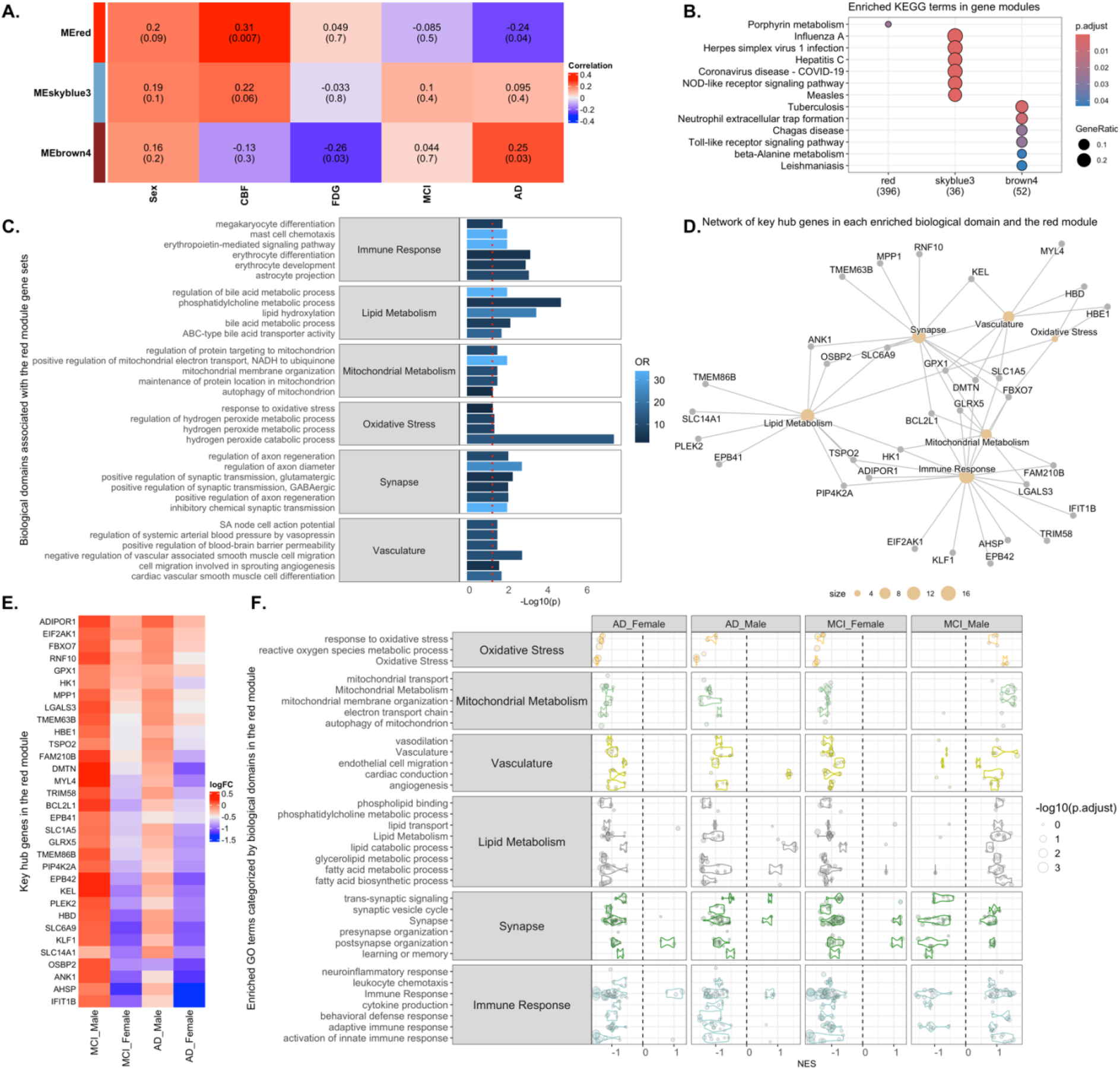
(**A**) WGCNA gene modules associated with average z-scores of CBF and FDG imaging data. The red gene module was associated with CBF (p<0.01) and AD status (p<0.05)), while the skyblue3 module was primarily associated with CBF (p < 0.1) and AD (p < 0.05), and the brown4 module was associated with FDG (p < 0.05) and AD status (p < 0.05). (**B**) Enriched KEGG pathways (adjusted p < 0.1) in the red, brown4, and skyblue3 module gene sets. (**C**) AD-related biological domain and resident GO terms enrichment analysis in the CBF driven red module gene set using Fisher exact test, with the top six enriched GO terms within each enriched bidomain. (**D**) Network of hub genes in each enriched biological domain and the red module. (**E**) Heatmap displaying the log fold change values of hub genes across the MCI and AD cases compared to sex-matched control cases. (**F**) Enriched GO terms categorized by AD biological domains in the red module gene sets for MCI male, MCI female, AD male, and AD female cases. AD, Alzheimer’s disease; CBF, cerebral blood flow; FDG, glucose uptake.

We further assessed the enrichment of AD biological domains^22^ within gene modules, and found that GO-terms associated with immune response, lipid and mitochondrial metabolism, oxidative stress, synapse, and vasculature biological domains were significantly enriched in CBF driven red modules (p<0.05; Figures 5.C-D/Table S7-S8). Next, we examined the changes in gene expression within the red module in male and female MCI and AD patients compared to sex-matched controls. We identified several genes showing distinct expression changes in MCI and AD patients, as well as genes with opposite expression changes in males and females (such as SLC14A1, PLEK2, and SLC6A9)(Figure 5.E/Table S9). Further, we ranked red module genes based on log fold change values calculated for each disease state and sex, and GSEA was performed. The GSEA revealed downregulation of multiple GO-terms related to immune response, lipid and mitochondrial metabolism, oxidative stress, synapse, and vasculature biological domains in female MCI, along with female and male AD patients. However, these same biological processes were upregulated in MCI male patients (Figure 5.F/Table S10).

Next, we investigated genes in the FDG associated with brown4 module. We observed that these genes were prominently enriched for GO-terms associated with immune response biological domain, such as “inflammatory response” and “microglial cell activation”, lipid metabolism biological domain, including “fatty acid omega-oxidation”, synapse, and vasculature biological domains (p<0.05; Figures 6.A-B/Table S7-S8). We identified several genes showing similar expression changes in the MCI and AD patients, such as SORL1, FCGR2A, and CDK19, as well as opposite gene expression changes in MCI and AD patients, such as STX3 and SULT1B1 (Figure 5.C/Table S9). The GSEA on these brown4 module genes revealed upregulation of multiple GO-term associated with immune response biological domain, such as “adaptive immune response” and “cytokine production”, proteostasis, and synapse biological domains in both MCI and AD cases (Figure 6.D/Table S11).

**Figure 6.**
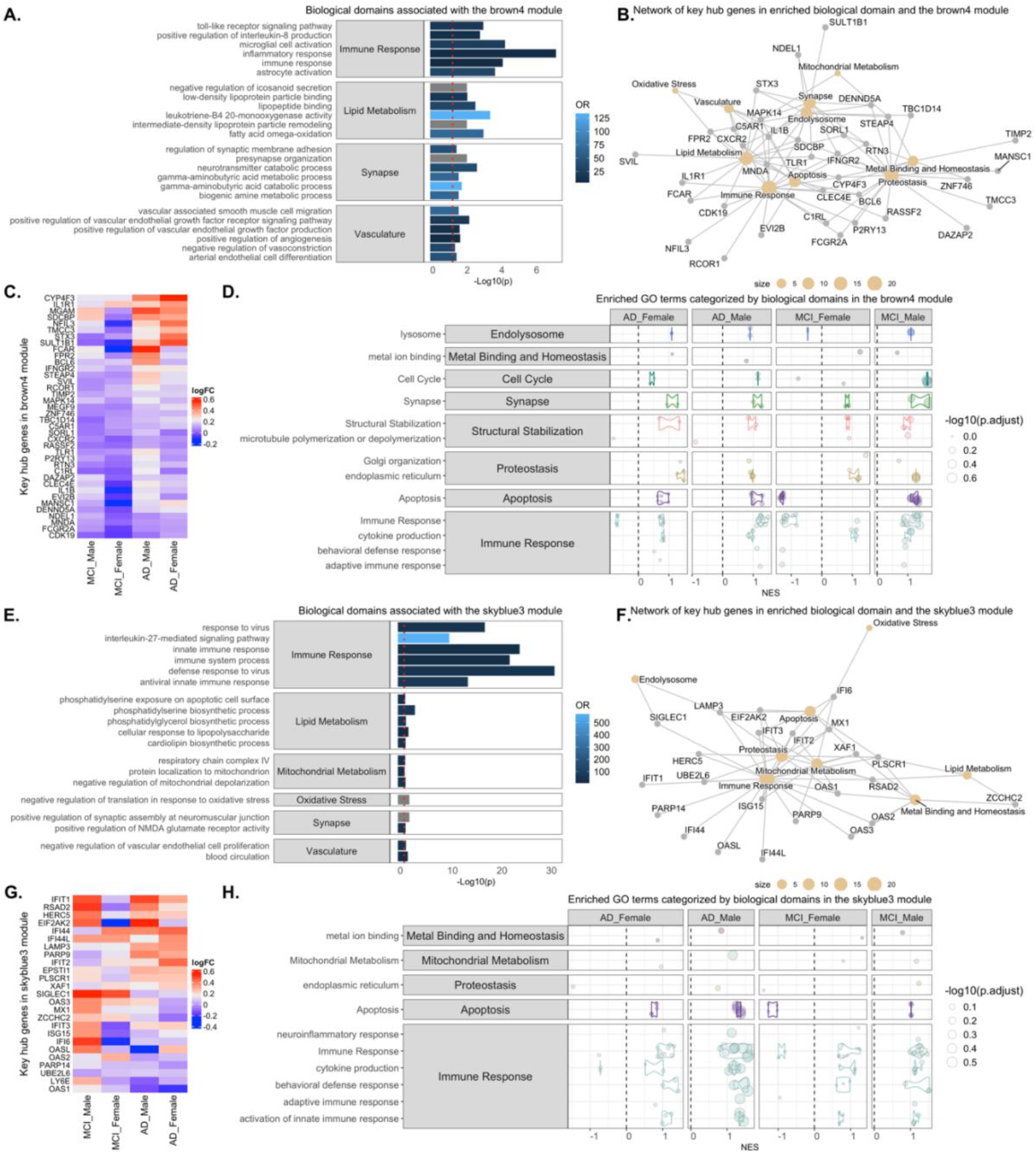
(**A**) AD-related biological domain and resident GO terms enrichment analysis in the FDG driven brown4 module gene set using Fisher exact test, with the top six enriched GO terms within each enriched bidomain. (**B**) Network of hub genes in each enriched biological domain and the brown4 module. (**C**) Heatmap displaying the log fold change values of hub genes in the brown4 module across the MCI and AD cases compared to sex-matched control cases. (**D**) Enriched GO terms categorized by AD biological domains in the brown4 module gene sets for MCI male, MCI female, AD male, and AD female cases. (**E**) AD-related biological domain and resident GO terms enrichment analysis in the FDG driven brown4 module gene set using Fisher exact test, with the top six enriched GO terms within each enriched bidomain. (**F**) Network of hub genes in each enriched biological domain and the brown4 module. (**G**) Heatmap displaying the log fold change values of hub genes in the brown4 module across the MCI and AD cases compared to sex-matched control cases. (**H**) Enriched GO terms categorized by AD biological domains in the brown4 module gene sets for MCI male, MCI female, AD male, and AD female cases.

Lastly, we investigated genes in the CBF associated skyblue3 module. We observed that these genes were prominently enriched for GO-terms associated with immune response biological domain, such as “innate immune response”, “interleukin-27-mediated signaling pathway”, and “immune system process”, (p<0.05; Figures 6.E-F/Table S7-S8). We also identified several genes showing distinct expression changes in the MCI and AD patients (Figure 6.G/Table S9). The GSEA on the skyblue3 module genes also revealed upregulation of multiple GO-term associated with Immune Response biological domain, such as “activation of innate immune response” and “cytokine production” in both MCI and AD cases (Figure 6.H/Table S12).

### 1.6 Relationship with Clinical Cognitive Assessment (CCA)

To align the MVD analysis of FDG and CBF SUVRs with CCA, we performed covariate analysis between brain regions and CCA test components for males and females (Figures S10-S11). To identify significant relationships, we performed Student’s t-tests (p<0.05) to identify slopes which were significantly different from zero. As anticipated, the bulk of the relationships were significant across both sexes for nearly all measures. For CBF SUVRs, correlation values ranged from [-0.33,0.36] for males (Figure S10.A) and [-0.27,0.36] for females (Figure S11.A), with means of 0.0082±0.0998 and 0.0201±0.0917 respectively, which suggests a limited connection between CBF changes and CCA scores. Comparatively, for FDG SUVRs, the values ranged from [-0.5,0.4] for males with a mean of - 0.0551±0.1528 (Figure S10.B) and from [-0.6,0.4] for females with a mean of -0.0484±0.1594 (Figure S11.B). Despite larger intervals, these correlations still indicate a weak association overall. Importantly, the data suggests that CCA may not have the fidelity to capture the metabolic and perfusion changes associated with disease progression. Moreover, this data indicates that the broad test categories used in the individual CCA’s might hinder accurate staging, since tests may struggle to differentiate between high and low metabolic/perfusion phases. Notably, regions showing positive, or negative, correlations do so consistently across all tests’ sections (Figures S10-S11).

## 2. Discussion

Our study confirmed that regional disease pathways map to MVD changes across AD progression. The disease pathways and MVD patterns observed, revealed the following: 1) in the early disease stage (CN⟶EMCI; Figure 2.A/2.E), there is an increase in perfusion (T1U) due to neuroinflammation and gliosis, which aligns with preclinical studies seen in mice.^8^ As the disease progresses into MCI (Figure 2.B/2.F), there is an increase in metabolism resultant from astrocytosis and energy shuttling to neurons which are impaired by significant oxidative stress (T2U). As more severe clinical symptoms appear (MCI⟶LMCI/AD; Figure 2.C/2.G), perfusion and metabolism increase to accommodate the increased regional demand; however, this increase in supply by astrocyte/neuron metabolic shuttling and increases in perfusion are finite, thus eroding the reserve capacity of the brain during the upregulation of neurons (PC). As demand exceeds reserve capacity to support brain function, during the latter stage of disease (AD; Figures 2.D/2.H), both metabolism and perfusion begin to fail, as the brain can only maintain the quasi-stable metabolic state for a limited time before the neurodegenerative damage becomes irreversible, due to cerebral metabolic and perfusion insufficiency (NMVF). This is consistent with our previous findings in rodent models of AD,^8,23^ and suggest that the regional MVD pattern (i.e. T1U⟶T2U⟶PC⟶NMVF) is generable across species. Moreover, our findings emphasize the delicate coordination between vascular, metabolic, and inflammatory processes across the disease spectrum, and reinforces the notion that a compensatory upregulation of perfusion and metabolism occurs in the early to mid-stages of AD;^5^ however, the precise inflection points vary by brain regions and individuals (Figures 3/4). Furthermore, our findings demonstrate that perfusion and metabolism eventually fail, are consistent with the “metabolic reprogramming theory”^7^ and the “energy failure hypothesis”,^16^ which suggesting that prolonged metabolic stress and neuroinflammation ultimately overwhelm the brain’s reserve capacity and causing restructuring of the brain networks in ADRD.

In addition, UMC analysis revealed that regions associated with memory, cognition, and motor functions exhibited different progression rates relative to CN, with early stages (EMCI/MCI) showing both T1U and T2U phenotypes, while late stages (LMCI/AD) exhibited PC and NMVF phenotypes. These findings align with prior findings that glial cells are activated to compensate for increased energetic demand,^19^ triggering metabolic reprogramming to support neuronal function under stress. Moreover, our data indicate that chronic neuroinflammation often precedes, and contributes, to vascular dysregulation,^17^ and are consistent with our previous reports in mouse models of AD.^8,23^ Importantly, the upregulation of neurons under oxidative stress and CBF insufficiency further compounds cerebral hypoxia, which has been linked to the accumulation of amyloid plaque and tau tangles as metabolic demand and vascular proliferation.^13,18^ Taken together, our findings reinforce the multifactorial nature of AD, illustrating how metabolic, vascular, and inflammatory processes converge to drive neuronal death and clinical symptom progression.

Importantly, UMC analysis revealed that brain regions follow MVD trajectory across the disease spectrum; however, brain regions follow different trajectories and unique rates (Figures 3/4). By contrast, some regions show little to no change in their trajectory angles and/or magnitudes. These results suggest that brain regions undergo structural and/or functional alterations that result in the reordering of the brain networks as a compensatory response to sustain the energetic load. The importance of the hippocampus, temporal gyrus, and amygdala in memory and learning, and the fact that these regions are the first to show both AD-related changes and significant neurovascular uncoupling, suggests that this diagnostic approach detects MVD at earlier stages than previous reports of single modality analyses.^2,3^ In addition, our study revealed significant sex and disease stage-dependent MVD in key brain regions, consistent with our previous work in AD mice,^8^ and suggest that trajectories for each brain region can be used as a unique signature to determine at-risk regions during disease progression.

Transcriptomics revealed changes in immune response, lipid and mitochondrial metabolism, oxidate stress, synapse, and vasculature biological domains for significant modules (Figure 5/6). For example, GO-terms associated with inflammatory response, microglial cell activation, fatty acid omega oxidation, cytokine production, proteostasis, and activation of innate immune responses showed prominent enrichment. This result aligns with the metabolic restructuring, energy deficits, and NVC dysregulation theories supporting that astrocytes play a role in disease progression by increasing energy production and supplying nutrients to neurons under stress conditions; however, this cannot be supported indefinitely.^7,8,12,14,16,22^ This aligns with our MVD results, which suggest that the brain undergoes a hypermetabolic phase in the early stages (EMCI/MCI) and then falls into a hypometabolic phase in the late stages (LMCI/AD)(Figures 2.A-D). Moreover, some genes showed similar expression changes (i.e., SORL1, FCGR2A, and CDK19); in contrast, others revealed opposite expression changes (i.e., STX3, SULT1B1); which indicates alterations in the regulation of the different genes influencing metabolism, signaling (i.e., cell stress, inflammatory), and vasculature restructuring. GSEA revealed downregulation for both sexes in AD but only for MCI females, while it was upregulated for MCI males (Figure 5.F). This aligns with our MVD analysis results (Figure 2) which shows that females progress faster through the disease spectrum.

Lastly, since CCA primarily focuses on memory, motor tasks, and recognition, suggests the reliability of these measures for predicting outcomes may be less reliable.^24^ Our findings demonstrate that the regions targeted by these tests exhibit accelerated progression through the MVD pattern (Figures 3/4), which suggests that brain connectivity networks between at-risk regions undergo alterations across the AD spectrum. To elucidate the MVD changes with clinical cognitive tests, correlation studies conducted between estimates of metabolism, perfusion and CCA demonstrated weak associations, suggesting limited predictability and non-linear associations between measures. Furthermore, our data indicates that CCA alone may not have the fidelity to stratify the subjects into finer disease stages, and that relying solely on CCA evaluations may overlook the subtle pathophysiological changes.^25^ Consequently, we propose that imaging biomarkers can be used to complement CCA for a more robust and accurate stratification of the disease stages, leading to a deeper insight into the disease etiology.

As with any research, this study has consideration and limitations that should be noted. First, we used z-scored SUVR values of the perfusion and metabolism measurements, as outlined in our previous preclinical study.^8^ This approach is applicable to any multi-modal context that can serve as surrogate biomarker to assess MVD, since scales/units are eliminated; however, the selection of an appropriate reference region/structure is strongly debated,^26,27^ and selection of a normalization factor may alter interpretations. Additionally, MVD and UMC plots are restricted in showing changes relative to a reference group, which must be clearly defined to permit accurate interpretation of the findings. Second, correlations among the various biomarkers only determine their similarities without considering their interrelationships. Third, our analysis was restricted to the study population level, so an inter-subject analysis is necessary to capture the range of variability within each disease stage group. Fourth, because the youngest age in our dataset was 55 years, data from younger individuals may be needed to characterize the complete MVD pattern. Finally, we obtained genetic data for only a subset of the study population. Hence, a more extensive dataset that contains genetics, CCA, and imaging data for all subjects would be desirable for verifying our findings.

In summary, our findings show that MVD of the at-risk brain regions depends on sex and disease stage. These changes were consistent with previously published work in preclinical mice studies, demonstrating the current approach is appropriate for detecting regions at-risk during the stages of ADRD progression. Moreover, this data suggests that these novel MVD patterns can provide a sensitive tool for early diagnosis of AD, which may improve patient monitoring, stratification, and therapeutic testing.

## 3. Online Methods

### 3.1 Dataset

Data used in the preparation of this article were obtained from the Alzheimer’s Disease Neuroimaging Initiative (ADNI) database (adni.loni.usc.edu). ADNI was launched in 2003 as a public-private partnership, led by Principal Investigator Michael W. Weiner, MD. The primary goal of ADNI has been to test whether serial MRI, PET, other biological markers, and clinical and neuropsychological assessment can be combined to measure the progression of MCI and early AD.^28^

Specifically, we mined data from ADNI phases 2 and 3 for subjects scanned with ^18^F-FDG PET (FDG), 3D T1-weighted, Fluid-Attenuated Inversion Recovery (FLAIR), and Arterial Spin Labeling (ASL) MRI within 180 days. A total of 403 cases were obtained, and split as follows: CN=95, EMCI=71, MCI=112, LMCI=42, and AD=83. Subjects ranged in age from 55 to 95 years, with a sex split of 55% males and 45% females. In addition, vitals data (i.e. age, height, weight), APOE genotype, and clinical cognitive assessment (CCA) results (ADAS, CDR, MoCA) for these subjects were obtained. Additionally, Genome-Wide Association Studies (GWAS) and gene expression profiles were available for a subpopulation (N=116: CN=77; MCI=18; AD=21) of these subjects.

### 3.2 Regional Metabolic and Vascular Analysis

To evaluate regional glycolysis, glucose uptake was measured via FDG as surrogate biomarker for glycolytic metabolism.^29^ FDG images were acquired and reconstructed according to standardized protocols developed by ADNI^28^ (Table S1). Regional perfusion was assessed by ASL MRI using standardized ADNI protocols^28^ with optimized parameters (Table S2). 3D T1-weighted and FLAIR images combined with 2D pulsed ASL (PASL) images from ADNI2/3 subjects, and 3D pseudo-continuous ASL (pCASL) and 3D PASL images from ADNI3 subjects were processed via ExploreASL^30^ yielding CBF^31^ maps. To permit direct comparison in a common reference space (i.e. Montreal Neurological Institute + Harvard-Oxford Cortical + subcortical(RRID:SCR_001476) + FSL Probabilistic Cerebellar Atlases; MNI152+), all images were co-registered using a 3D deformable registration developed in our lab.^32^

Post-registration, mean intensities for 59 unilateral regions were extracted for both FDG and CBF images ratioed to whole brain activity, and converted to z-scores relative to the reference population. To estimate the extent of neurovascular uncoupling, standardized uptake value ratios (SUVRs) relative to the whole brain^33^ were computed using:

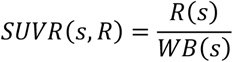

where, *s, R*, and *WB* denote the subject, mean value of the region/volume of interest for subject *s*, and whole brain mean value for subject *s*, respectivley. For each regional SUVR, values were converted to z-scores relative to the reference population using:

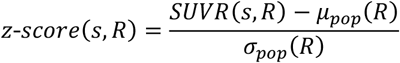

where, *s, R, µ*_*pop*_, and *σ*_*pop*_ denote subject, region/volume of interest, mean SUVR of the reference population, and standard deviation of the reference population, respectively.

To permit tracking of individual regions across the disease spectrum, and to facilitate tracking of region trajectories, Cartesian coordinates were transformed into Polar coordinates to visualize the disease progression while retaining NVC state. The distance from the origin indicates the magnitude of change and the angle determines the deviation of the change (Figure 1.C). Data was projected onto 2D Cartesian space, where the x-axis represents the z-scores derived from CBF maps, and the y-axis the z-score derived from FDG (Figure 1.B). The transformation into Polar coordinate is given by:

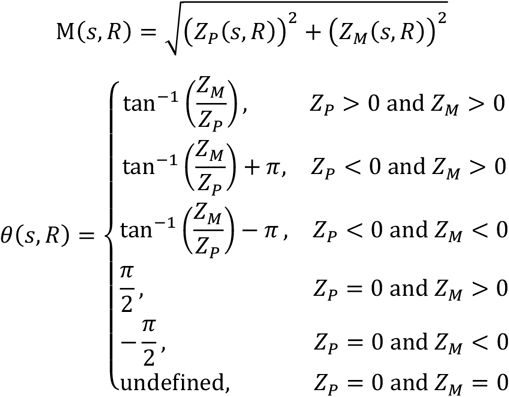

where, *s, R, Z*_*p*_, *Z*_*M*_, *M* and *θ* denote the subject, the region/volume of interest for subject *s*, the z-score changes in CBF, the z-score change in glucose uptake, the magnitude of the change, and the direction of the change in the polar space, respectively.

### 3.3 Transcriptomics Analysis

#### 3.3.1 Weighted gene co-expression network analysis

Weighted gene co-expression network analysis (WGCNA)^34^ was conducted to identify clusters of genes with similar expression pattern using log-normalized gene expression data from blood samples of ADNI participants. A default unsigned network type was used, with a soft thresholding power of 6 to satisfy the scale-free topology criterion in the pickSoftThreshold function.^21^ We set the minimum modules size as 30, and merged modules whose correlation coefficients were greater than 0.75 (mergeCutHeight = 0.25). Each module was summarized by its module eigengene (ME), the first principal component of the gene expression profiles within a module. We also identified key genes (hub genes) within modules that have a module membership (kME) greater than 0.75. Module membership indicates the degree to which a gene is part of a module. High module membership values mean a gene’s expression profile is highly correlated with the module eigengene, suggesting that the gene is a central or hub gene within that module. Additionally, we calculated the Pearson correlation coefficient between the modules and various participant characteristics, including sex, AD and MCI status, CBF, and FDG measures. To visualize the relationships between gene expression data and external traits, we employed the ComplexHeatmap package in R.

#### 3.3.2 Functional enrichment analysis

Functional annotations and enrichment analyses were performed using the R Bioconductor package clusterProfiler,^35^ with Gene Ontology term and KEGG pathway enrichment analyses performed using enrichGO and enrichKEGG, respectively. The compareCluster function was used to compare enriched functional categories of each gene module. The significance threshold for all enrichment analyses was set to 0.05 using Benjamini-Hochberg adjusted p-values.

#### 3.3.3 Enrichment of AD biological domains

Gene functional enrichment analyses are informative but sometimes it is unclear how the enriched terms relate to the biology of AD. Cary et al. ^22^ developed 19 biological domains that capture the AD-associated endophenotypes and defined them using an exhaustive set of Gene Ontology (GO) terms, with the intent to keep each domain siloed in a biologically coherent fashion. First, we performed Fisher’s exact tests to determine the significance of the overlap between genes in each AD biological domain and resident GO terms in WGCNA module gene sets. The cnetplot function in the enrichplot package was used for visualizing gene-concept networks based on enrichment analysis results. We also measured the log fold change values of genes for MCI and AD cases compared to sex-matched controls, calculated using the lmFit function from the limma package^36^ Genes in each module exhibit different expression changes in the MCI male, MCI female, AD male, and AD female patients compared to controls. Therefore, we performed Gene Set Enrichment Analysis (GSEA)^37^ using the same genes sorted in descending order based on log fold change values as input within modules, separately for MCI male, MCI female, AD male, and AD female participants. GSEA was performed using the gseGO function from the clusterProfiler R package^35^ and the results were then categorized into biological domains and sub-domains based on the GO ID of enriched terms. The displayed results include the normalized enrichment score (NES) as well as the Benjamini-Hochberg corrected p value (p adj) for GO terms annotated to each biological domain.

### 3.4 Statistical Analysis

Statistical analysis of at-risk regions was evaluated using Student’s t-test (p<0.05), without Bonferroni correction since regional independence was maintained, where neuro-metabolic-vascular uncoupling phenotype was determined by the Cartesian location of glycolytic metabolism and CBF z-scores. Significant regions were then projected onto MNI152+ object maps to provide spatial localization, where the sign of the z-score was applied to the p-value to indicate directionality of change (i.e. 0<z increasing, z<0 decreasing). In addition, the correlation between CBF and FDG SUVR values of the brain regions, and CCA were aligned to evaluate their relationship across the disease spectrum, and Student’s t-test (p<0.05) used to determine their significance.

## Supporting information

Supplemental Sections

## Acknowledgements

Data collection and sharing for this project was funded by the Alzheimer’s Disease Neuroimaging Initiative (ADNI) (National Institutes of Health Grant U01 AG024904) and DOD ADNI (Department of Defense award number W81XWH-12-2-0012). ADNI is funded by the National Institute on Aging, the National Institute of Biomedical Imaging and Bioengineering, and through generous contributions from the following: AbbVie, Alzheimer’s Association; Alzheimer’s Drug Discovery Foundation; Araclon Biotech; BioClinica, Inc.; Biogen; Bristol-Myers Squibb Company; CereSpir, Inc.; Cogstate; Eisai Inc.; Elan Pharmaceuticals, Inc.; Eli Lilly and Company; EuroImmun; F. Hoffmann-La Roche Ltd and its affiliated company Genentech, Inc.; Fujirebio; GE Healthcare; IXICO Ltd.; Janssen Alzheimer Immunotherapy Research & Development, LLC.; Johnson & Johnson Pharmaceutical Research & Development LLC.; Lumosity; Lundbeck; Merck & Co., Inc.; Meso Scale Diagnostics, LLC.; NeuroRx Research; Neurotrack Technologies; Novartis Pharmaceuticals Corporation; Pfizer Inc.; Piramal Imaging; Servier; Takeda Pharmaceutical Company; and Transition Therapeutics. The Canadian Institutes of Health Research is providing funds to support ADNI clinical sites in Canada. Private sector contributions are facilitated by the Foundation for the National Institutes of Health (www.fnih.org). The grantee organization is the Northern California Institute for Research and Education, and the study is coordinated by the Alzheimer’s Therapeutic Research Institute at the University of Southern California. ADNI data are disseminated by the Laboratory for Neuro Imaging at the University of Southern California. We thank Dorsey Faught from MIM Software Inc who helped with the use and development of workflows in MIM.

## Author’s Contributions

JAKCC contributed to the study design, data collection and management, data analysis and interpretation, data quality assessment, literature review, and manuscript drafting. SAC participated in study design, data collection and management, data analysis, and quality assessment. RSP provided transcriptomics analysis and interpretation, and helped write the manuscript. OS did literature search for functional analysis of the brain regions. PS provided statistical and image analysis advice, participated in data interpretation, and helped write the manuscript. PRT contributed to the study design, data analysis and interpretation, and helped write the manuscript. JAKCC, PRT, and SAC had access to all raw data. RSP had access to raw genetic data. JAKCC, PRT, and SAC verified the data and results. All authors helped revise the manuscript. All authors had final responsibility for the decision to submit for publication.

## Data Sharing

The data that support the findings of this study were obtained from the Alzheimer’s Disease Neuroimaging Initiative (ADNI), which is available from the ADNI database (https://adni.loni.usc.edu) upon registration and compliance with the data use agreement.

## Ethics Committee Approval

Ethics approval was not required for this study.

## References

1. 2024 Alzheimer’s disease facts and figures. Alzheimers Dement. 2024;20(5):3708–821.

2. Canal-Garcia A, Vereb D, Mijalkov M, Westman E, Volpe G, Pereira JB, et al. Dynamic multilayer functional connectivity detects preclinical and clinical Alzheimer’s disease. Cereb Cortex. 2024;34(2).

3. Warren SL, Moustafa AA. Functional magnetic resonance imaging, deep learning, and Alzheimer’s disease: A systematic review. J Neuroimaging. 2023;33(1):5–18.

4. Stackhouse TL, Mishra A. Neurovascular coupling in development and disease: focus on astrocytes. Frontiers in cell and developmental biology. 2021;9:702832.

5. Apatiga-Perez R, Soto-Rojas LO, Campa-Cordoba BB, Luna-Viramontes NI, Cuevas E, Villanueva-Fierro I, et al. Neurovascular dysfunction and vascular amyloid accumulation as early events in Alzheimer’s disease. Metab Brain Dis. 2022;37(1):39–50.

6. Fontana IC, Scarpa M, Malarte ML, Rocha FM, Auselle-Bosch S, Bluma M, et al. Astrocyte Signature in Alzheimer’s Disease Continuum through a Multi-PET Tracer Imaging Perspective. Cells. 2023;12(11).

7. Demetrius LA, Eckert A, Grimm A. Sex differences in Alzheimer’s disease: metabolic reprogramming and therapeutic intervention. Trends Endocrinol Metab. 2021;32(12):963–79.

8. Onos KD, Lin PB, Pandey RS, Persohn SA, Burton CP, Miner EW, et al. Assessment of neurovascular uncoupling: APOE status is a key driver of early metabolic and vascular dysfunction. Alzheimers Dement. 2024;20(7):4951–69.

9. Kazemeini S, Nadeem-Tariq A, Shih R, Rafanan J, Ghani N, Vida TA. From Plaques to Pathways in Alzheimer’s Disease: The Mitochondrial-Neurovascular-Metabolic Hypothesis. Int J Mol Sci. 2024;25(21).

10. Heneka MT, van der Flier WM, Jessen F, Hoozemanns J, Thal DR, Boche D, et al. Neuroinflammation in Alzheimer disease. Nat Rev Immunol. 2024.

11. Shen L, Kim S, Risacher SL, Nho K, Swaminathan S, West JD, et al. Whole genome association study of brain-wide imaging phenotypes for identifying quantitative trait loci in MCI and AD: A study of the ADNI cohort. Neuroimage. 2010;53(3):1051–63.

12. Xiang X, Wind K, Wiedemann T, Blume T, Shi Y, Briel N, et al. Microglial activation states drive glucose uptake and FDG-PET alterations in neurodegenerative diseases. Sci Transl Med. 2021;13(615):eabe5640.

13. Bai R, Guo J, Ye XY, Xie Y, Xie T. Oxidative stress: The core pathogenesis and mechanism of Alzheimer’s disease. Ageing Res Rev. 2022;77:101619.

14. Brandebura AN, Paumier A, Onur TS, Allen NJ. Astrocyte contribution to dysfunction, risk and progression in neurodegenerative disorders. Nat Rev Neurosci. 2023;24(1):23–39.

15. Wang Z, Bovik AC, Sheikh HR, Simoncelli EP. Image quality assessment: from error visibility to structural similarity. IEEE transactions on image processing. 2004;13(4):600–12.

16. Yassine HN, Solomon V, Thakral A, Sheikh-Bahaei N, Chui HC, Braskie MN, et al. Brain energy failure in dementia syndromes: Opportunities and challenges for glucagon-like peptide-1 receptor agonists. Alzheimers Dement. 2022;18(3):478–97.

17. Al-Ghraiybah NF, Wang J, Alkhalifa AE, Roberts AB, Raj R, Yang E, et al. Glial Cell-Mediated Neuroinflammation in Alzheimer’s Disease. Int J Mol Sci. 2022;23(18).

18. Vyas J, Raytthatha N, Prajapati BG. Chapter 6 - Amyloid cascade hypothesis, tau synthesis, and role of oxidative stress in AD. In: Prajapati BG, Chellappan DK, Kendre PN, editors. Alzheimer’s Disease and Advanced Drug Delivery Strategies: Academic Press; 2024. p. 73–92.

19. Fan Z, Okello AA, Brooks DJ, Edison P. Longitudinal influence of microglial activation and amyloid on neuronal function in Alzheimer’s disease. Brain. 2015;138(Pt 12):3685–98.

20. Tsartsalis S, Sleven H, Fancy N, Wessely F, Smith AM, Willumsen N, et al. A single nuclear transcriptomic characterisation of mechanisms responsible for impaired angiogenesis and blood-brain barrier function in Alzheimer’s disease. Nat Commun. 2024;15(1):2243.

21. Langfelder P, Horvath S. WGCNA: an R package for weighted correlation network analysis. BMC Bioinformatics. 2008;9:559.

22. Cary GA, Wiley JC, Gockley J, Keegan S, Amirtha Ganesh SS, Heath L, et al. Genetic and multi-omic risk assessment of Alzheimer’s disease implicates core associated biological domains. Alzheimers Dement (N Y). 2024;10(2):e12461.

23. Kotredes KP, Pandey RS, Persohn S, Elderidge K, Burton CP, Miner EW, et al. Characterizing molecular and synaptic signatures in mouse models of late-onset Alzheimer’s disease independent of amyloid and tau pathology. Alzheimers Dement. 2024;20(6):4126–46.

24. Clarke A, Ashe C, Jenkinson J, Rowe O, A DNI, Hyland P, et al. Predicting conversion of patients with Mild Cognitive Impairment to Alzheimer’s disease using bedside cognitive assessments. J Clin Exp Neuropsychol. 2022;44(10):703–12.

25. Vemuri P, Wiste HJ, Weigand SD, Shaw LM, Trojanowski JQ, Weiner MW, et al. MRI and CSF biomarkers in normal, MCI, and AD subjects: diagnostic discrimination and cognitive correlations. Neurology. 2009;73(4):287–93.

26. Zhao Z, Wang J, Wang Y, Liu X, He K, Guo Q, et al. 18F-AV45 PET and MRI Reveal the Influencing Factors of Alzheimer’s Disease Biomarkers in Subjective Cognitive Decline Population. J Alzheimers Dis. 2023;93(2):585–94.

27. Dang C, Wang Y, Li Q, Lu Y. Neuroimaging modalities in the detection of Alzheimer’s disease-associated biomarkers. Psychoradiology. 2023;3:kkad009.

28. Mueller SG, Weiner MW, Thal LJ, Petersen RC, Jack C, Jagust W, et al. The Alzheimer’s disease neuroimaging initiative. Neuroimaging Clin N Am. 2005;15(4):869-77, xi-xii.

29. Perovnik M, Tang CC, Namias M, Eidelberg D, Alzheimer’s Disease Neuroimaging I. Longitudinal changes in metabolic network activity in early Alzheimer’s disease. Alzheimers Dement. 2023;19(9):4061–72.

30. Mutsaerts H, Petr J, Groot P, Vandemaele P, Ingala S, Robertson AD, et al. ExploreASL: An image processing pipeline for multi-center ASL perfusion MRI studies. Neuroimage. 2020;219:117031.

31. Dijsselhof M, James S-N, Lorenzini L, Collij L, Thomas D, Scott C, et al. Cerebral blood flow as intermediary between cardiovascular and cerebrovascular health: results from the Insight46 study. Cerebral Circulation - Cognition and Behavior. 2024;6:100261.

32. Chong Chie JAK, Persohn SC, Miner EW, Burton CP, Salama P, Territo PR, editors. Total Variation Based 2D Image Registration of Post-Mortem Mouse Brain Images. 2024 IEEE International Symposium on Biomedical Imaging (ISBI); 2024: IEEE.

33. Nugent S, Croteau E, Potvin O, Castellano CA, Dieumegarde L, Cunnane SC, et al. Selection of the optimal intensity normalization region for FDG-PET studies of normal aging and Alzheimer’s disease. Sci Rep. 2020;10(1):9261.

34. Zhang B, Horvath S. A general framework for weighted gene co-expression network analysis. Stat Appl Genet Mol Biol. 2005;4:Article17.

35. Yu G, Wang LG, Han Y, He QY. clusterProfiler: an R package for comparing biological themes among gene clusters. OMICS. 2012;16(5):284–7.

36. Ritchie ME, Phipson B, Wu D, Hu Y, Law CW, Shi W, et al. limma powers differential expression analyses for RNA-sequencing and microarray studies. Nucleic Acids Res. 2015;43(7):e47.

37. Subramanian A, Tamayo P, Mootha VK, Mukherjee S, Ebert BL, Gillette MA, et al. Gene set enrichment analysis: a knowledge-based approach for interpreting genome-wide expression profiles. Proc Natl Acad Sci U S A. 2005;102(43):15545–50.

