## Supplemental Sections for "Neuro-Metabolic and Vascular Dysfunction as an Early Diagnostic for Alzheimer’s Disease and Related Dementias"

### SUPPLEMENTAL MATERIAL

#### A. SUPPLEMENTAL TABLES

**Supplemental Table S1**. FDG Acquisition and Reconstruction Protocol used in ADNI 2/3 cycles.

| Phase | Manufacturer | Model | Radiotracer (mCi) | Scan Start Time (min) | Acquisition Mode | Duration and Framing | Reconstruction Method | Method Parameters | Grid | FOV | Slice Thickness (mm) | Smoothing |
|---|---|---|---|---|---|---|---|---|---|---|---|---|
| ADNI2 | GE | Discovery STE / VCT | 4.5 – 5.5 | 30 | N/A | 30 min, six x 5min frames | 3D IR | 4 iterations; 20 subsets | 128 x 128 | 256.00 | 3.270 | None |
| | | Discovery ST | | | | | | 4 iterations; 21 subsets | | | | |
| | | Discovery RX | | | | | | | | | | |
| | | Discovery LS | | | | | 3D FORE IR | | | | 4.250 | |
| | | Advance | | | | | | | | | | |
| | Phillips | Gemini TF | | | N/A | 30 min, six x 5min frames | 3D LOR-RAMLA | N/A | 128 x 128 | 256.00 | 2.000 | Sharp |
| | | Gemini / Gemini GXL | | | | | | | | | | |
| | | Allegro | | | | | | | | | | |
| | Siemens | BioGraph mCT | | | LIST-MODE | 30 min, six x 5min frames | OSEM3D | 4 iterations; 12 subsets | 400 x 400 | 407.20 | 2.027 | None |
| | | | | | No LIST-MODE | Two 15min scans | | | | | | |
| | | BioGraph TruePoint (1093/1094) | | | LIST-MODE | 30 min, six x 5min frames | OSEM2D | 4 iterations; 14 subsets | 336 x 336 | 341.04 | 2.027 | None |
| | | | | | No LIST-MODE | Two 15min scans | | | | | | |
| | | | | | LIST-MODE | 30 min, six x 5min frames | OSEM2D | 4 iterations; 16 subsets | 336 x 336 | 341.04 | 2.027 | None |
| | | | | | No LIST-MODE | Two 15min scans | | | | | | |
| | | BioGraph HiRes (1080) | | | LIST-MODE | 30 min, six x 5min frames | OSEM2D | 4 iterations; 14 subsets | 168 x 168 | 341.21 | 2.000 | None |
| | | | | | No LIST-MODE | Two 15min scans | | | | | | |
| ADNI3 | GE | Discovery 600, 610, 690, 710 | 5 ± 10% | 30 | N/A | 30 min, six x 5min frames | 3D IR | 4 iterations; 24 subsets | 192 x 192 | 256.00 | 3.270 | None |
| | | Discovery STE | | | | | | 4 iterations; 20 subsets | 128 x 128 | | | |
| | | | | | | | | 4 iterations; 21 subsets | | | | |
| | | Discovery ST | | | | | | | | | | |
| | | Discovery LS | | | | | 3D FORE IR | | | | 4.250 | |
| | | Advance | | | | | | | | | | |
| | Phillips | Ingenuity TF | | | N/A | 30 min, six x 5min frames | BLOB-OS-TF | N/A | 128 x 128 | 256.00 | 2.000 | Sharp |
| | | Gemini TF | | | | | 3D LOR-RAMLA | | | | | |
| | | Gemini / Gemini GXL | | | | | | | | | | |
| | | Allegro | | | | | | | | | | |
| | Siemens | BioGraph mCT | | | LIST-MODE | 30 min, six x 5min frames | OSEM3D | 4 iterations; 24 subsets | 400 x 400 | 407.20 | 2.027 | None |
| | | | | | No LIST-MODE | Two 15min scans | | | | | | |
| | | BioGraph TruePoint (1093/1094) | | | LIST-MODE | 30 min, six x 5min frames | OSEM3D | 4 iterations; 21 subsets | 336 x 336 | 341.04 | 2.027 | None |
| | | | | | No LIST-MODE | Two 15min scans | | | | | | |
| | | BioGraph HiRes (1080) | | | LIST-MODE | 30 min, six x 5min frames | OSEM2D | 4 iterations; 16 subsets | 168 x 168 | 341.04 | 2.000 | None |
| | | | | | No LIST-MODE | Two 15min scans | | | | | | |

**Supplemental Table S2**. Single PLD ASL MRI Acquisition used in ADNI 2/3 cycles.

| | **Siemens 2D PASL (ADNI 2/3)** | **Siemens 3D PASL (ADNI 3)** | **GE 3D pCASL (ADNI 3)** |
|---|---|---|---|
| Software Versions | 20VB17 | Prisma 20180612<br>Prisma D13<br>Prisma VE11C<br>Skyra E11<br>Skyra VE11<br>Magento Vida-XT | 25x<br>Widebore 25x |
| Repetition Time (TR) | 3400 ms | 4000 ms | 4888 ms |
| Echo Time (TE) | 12 ms | 20.26 – 21.80 ms | 10.528 |
| Field of View | 256 x 256 $mm^2$ | 240 x 240 $mm^2$ | 240 x 240 $mm^2$ |
| Acquisition Matrix | 64 x 64 | 64 x 64 | 128 x 128 |
| Reconstruction Matrix | 64 x 64 | 128 x 128 | 128 x 128 |
| Slice Thickness | 4 mm | 4.5 mm | 4 mm |
| Tag Thickness | 100 mm | N/A | N/A |
| Number of Slices | 24 | 32 | 40 |
| Bandwidth | 2368 Hz/pix | 2442 – 2604 Hz/pix | 976.6 Hz/pix |
| Background Suppression | Yes | Yes | Yes |
| M0 Available | Yes | No | Yes |
| Control-Tag Pairs | Yes, 52 | Yes, 10 | Yes, 40 |
| Mode/Readout | PICORE<br>Q2TIPS | GRASE | Stack of spirals |
| Bolus Duration | 700 ms | 800 ms | 1800 ms |
| Inversion Time / PLD | 1200 ms | 2000 ms | 2025 ms |
| Number of Repeats | 54 | 10 | 3 |
| Acquisition Time | 6:02 min | 5:24 – 8:04 min | 6 min |

**Supplemental Table S3**. Complete Demographics Stats

| | Count (n) | Age (years) | APOE (n) (ε2*, ε33, ε4*) | BMI | | | | |
|---|---|---|---|---|---|---|---|---|
| | | | | | Male | (n) | Female | (n) |
| CN | M 52 | M 76.75 +- 7.08 | M 7, 28, 17 | Underweight | N/A | 0 | 18.37 ± 0.00 | 1 |
| | | | | Normal | 23.63 ± 1.26 | 16 | 23.05 ± 1.89 | 17 |
| | | | | Overweight | 26.75 ± 1.11 | 31 | 27.74 ± 1.53 | 13 |
| | F 43 | F 74.86 ± 6.22 | F 4, 27, 12 | Obesity I | 31.51 ± 0.90 | 4 | 32.14 ± 1.61 | 10 |
| | | | | Obesity II | 36.15 ± 0.00 | 1 | N/A | 0 |
| | | | | Extreme Obesity | N/A | 0 | 49.76 ± 2.02 | 2 |
| EMCI | M 44 | M 72.08 ± 6.75 | M 5, 22, 17 | Underweight | N/A | 0 | 18.39 ± 0.00 | 1 |
| | | | | Normal | 22.89 ± 1.40 | 14 | 21.37 ± 2.02 | 9 |
| | | | | Overweight | 26.81 ± 1.30 | 21 | 27.42 ± 1.51 | 13 |
| | F 27 | F 69.75 ± 6.21 | F 3, 14, 10 | Obesity I | 32.20 ± 1.62 | 8 | 32.58 ± 0.61 | 4 |
| | | | | Obesity II | 37.34 ± 0.00 | 1 | N/A | 0 |
| | | | | Extreme Obesity | N/A | 0 | N/A | 0 |
| MCI | M 61 | M 74.52 ± 6.45 | M 3, 37, 21 | Underweight | N/A | 0 | 17.83 ± 0.00 | 1 |
| | | | | Normal | 23.32 ± 1.41 | 16 | 22.69 ± 1.76 | 19 |
| | | | | Overweight | 27.27 ± 1.48 | 34 | 27.75 ± 1.60 | 19 |
| | F 51 | F 71.16 ± 8.18 | F 3, 23, 25 | Obesity I | 31.40 ± 1.60 | 10 | 32.01 ± 1.45 | 9 |
| | | | | Obesity II | N/A | 0 | 36.25 ± 0.00 | 1 |
| | | | | Extreme Obesity | 43.20 ± 0.00 | 1 | 40.83 ± 0.97 | 2 |
| LMCI | M 17 | M 72.25 ± 7.62 | M 2, 8, 7 | Underweight | N/A | 0 | N/A | 0 |
| | | | | Normal | 24.21 ± 0.47 | 3 | 21.93 ± 0.84 | 9 |
| | | | | Overweight | 27.79 ± 0.98 | 10 | 27.07 ± 1.27 | 11 |
| | F 25 | F 72.77 ± 6.27 | F 1, 9, 15 | Obesity I | 31.37 ± 1.45 | 4 | 31.73 ± 1.28 | 4 |
| | | | | Obesity II | N/A | 0 | N/A | 0 |
| | | | | Extreme Obesity | N/A | 0 | 45.83 ± 0.00 | 1 |
| AD | M 48 | M 74.36 ± 8.78 | M 1, 16, 31 | Underweight | 18.09 ± 0.49 | 2 | 16.75 ± 0.00 | 1 |
| | | | | Normal | 22.44 ± 1.91 | 16 | 22.29 ± 1.84 | 19 |
| | | | | Overweight | 27.09 ± 1.66 | 26 | 27.77 ± 1.69 | 10 |
| | F 35 | F 73.98 ± 7.70 | F 0, 12, 23 | Obesity I | 30.90 ± 1.02 | 2 | 32.60 ± 1.18 | 4 |
| | | | | Obesity II | 37.58 ± 0.00 | 1 | 38.34 ± 0.00 | 1 |
| | | | | Extreme Obesity | 40.50 ± 0.00 | 1 | N/A | 0 |

**Table S4 – Complete Brain Regions Indices and Names.**

| Regions Index | | | |
|---|---|---|---|
| 01 - Frontal Pole | 16 - Inferior Temp Gyrus temporooccipit | 31 - Precuneus Cortex | 46 - Planum Temporale |
| 02 - Insular Cortex | 17 - Postcentral Gyrus | 32 - Cuneal Cortex | 47 - Supracalcarine Cortex |
| 03 - Superior Frontal Gyrus | 18 - Sup Parietal Lobule | 33 - Frontal Orbital Cortex | 48 - Occipital Pole |
| 04 - Mid Frontal Gyrus | 19 - Supramarginal Gyrus ant | 34 - Parahippocampal Gyrus ant | 49 - Cerebral White Matter |
| 05 - Inferior Frontal Gyrus pars trian | 20 - Supramarginal Gyrus post | 35 - Parahippocampal Gyrus post | 50 - Lateral Ventricle |
| 06 - Inferior Frontal Gyrus pars operc | 21 - Angular Gyrus | 36 - Lingual Gyrus | 51 - Thalamus |
| 07 - Precentral Gyrus | 22 - Lateral Occipital Cortex Sup | 37 - Temp Fusiform Cortex ant | 52 - Caudate |
| 08 - Temporal Pole | 23 - Lateral Occipital Cortex Inf | 38 - Temp Fusiform Cortex post | 53 - Putamen |
| 09 - Superior Temporal Gyrus ant | 24 - Intracalcarine Cortex | 39 - Temporal Occipital Fusiform Cortex | 54 - Pallidum |
| 10 - Superior Temporal Gyrus post | 25 - Frontal Medial Cortex | 40 - Occipital Fusiform Gyrus | 55 - Brain Stem |
| 11 - Mid Temp Gyrus ant | 26 - Juxtapositional Lobule Cortex | 41 - Frontal Operculum Cortex | 56 - Hippocampus |
| 12 - Mid Temp Gyrus post | 27 - Subcallosal Cortex | 42 - Central Opercular Cortex | 57 - Amygdala |
| 13 - Mid Temp Gyrus temporoocc | 28 - Paracingulate Gyrus | 43 - Parietal Operculum Cortex | 58 - Accumbens |
| 14 - Inferior Temp Gyrus ant | 29 - Cingulate Gyrus ant | 44 - Planum Polare | 59 - Cerebellum |
| 15 - Inferior Temp Gyrus post | 30 - Cingulate Gyrus post | 45 - Heschls Gyrus H1 H2 | |

**Supplemental Tables Captions from S5 to S12**

**Supplemental Table S5** – WGCNA modules of co-expressed genes: Genes in each of 28 modules of co-expressed genes identified through WGCNA of blood transcriptomic data from ADNI cohort.

**Supplemental Table S6** – Enriched KEGG pathways in the red, brown4, and skyblue4 gene modules.

**Supplemental Table S7** – Enrichment of AD biological domains in modules. Enrichment of AD biological domains and resident GO-terms in the red, brown4, and skyblue3 modules.

**Supplemental Table S8** – Genes overlapping in each AD biological domain and within the red, brown4, and skyblue3 modules.

**Supplemental Table S9** – logFold change values of genes in the MCI male, MCI female, AD male, and AD female cases compared to sex-matched controls.

**Supplemental Table S10** – Gene Set Enrichment Analysis results in the red module. Enriched GO terms in the red module genes ranked by log fold change values in MCI male, MCI female, AD male, and AD females. Enriched GO-terms were further categorized by AD biological domains.

**Supplemental Table S11** – Gene Set Enrichment Analysis results in the brown4 module. Enriched GO terms in the brown4 module genes ranked by log fold change values in MCI male, MCI female, AD male, and AD females. Enriched GO-terms were further categorized by AD biological domains.

**Supplemental Table S12** – Gene Set Enrichment Analysis results in the skyblue3 module. Enriched GO terms in the skyblue3 module genes ranked by log fold change values in MCI male, MCI female, AD male, and AD females. Enriched GO-terms were further categorized by AD biological domains.

**B. SUPPLEMENTAL FIGURES**

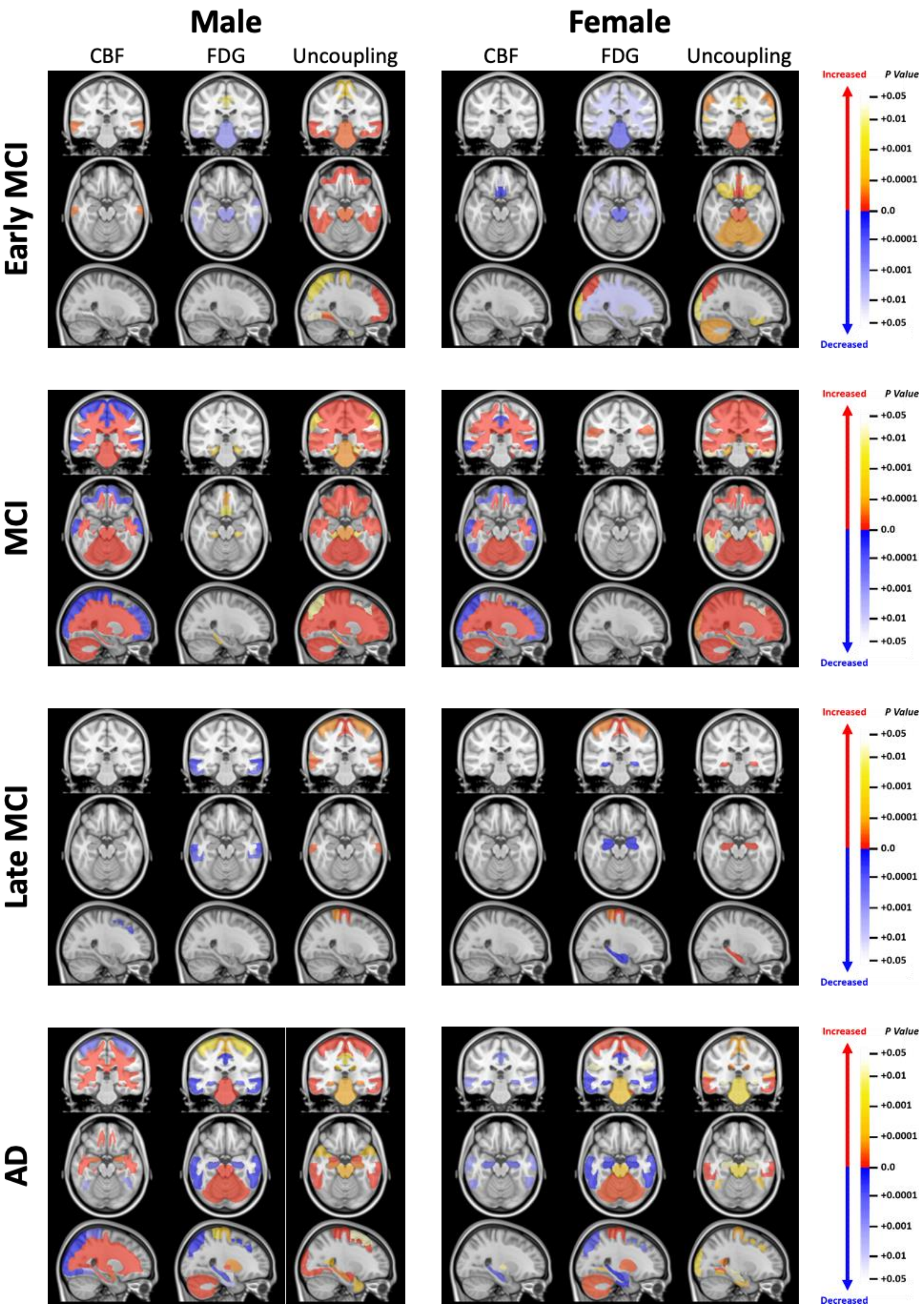


**Supplemental Figure S1 – Spatially Localized Significant Regions**
Significant regions (p<0.05) were projected onto MNI152+ object maps to determine their spatial localization. The sign of the z-score was then applied to the p-value to indicate the directionality of the change (i.e., a positive z-score indicates an increasing p-value, while a negative z-score indicates a decreasing p-value).

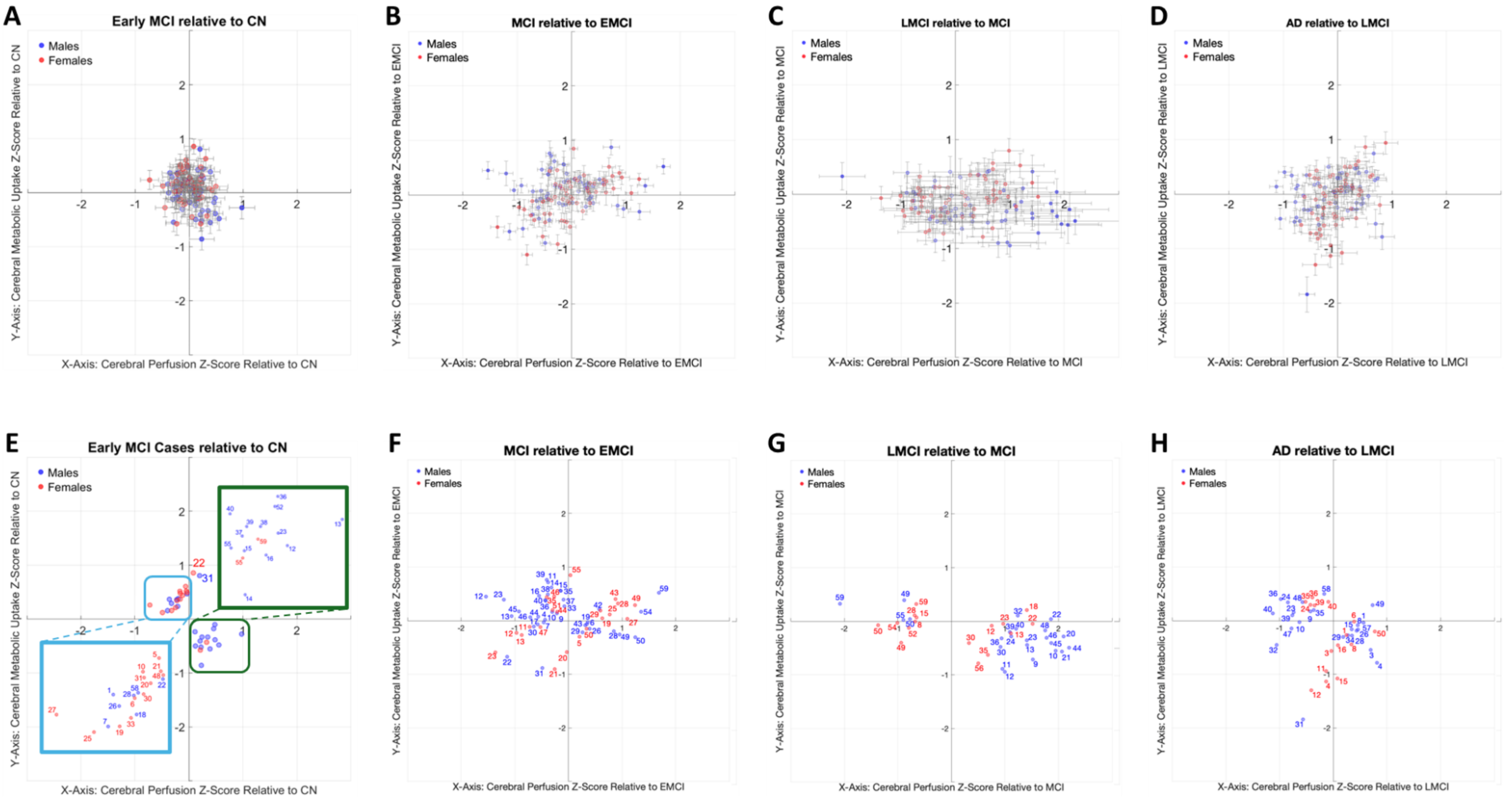


**Supplemental Figure S2 – Assessment of Neurovascular Coupling and Uncoupling of EMCI, MCI, LMCI, and AD Relative to the Previous Disease Stage.** To identify inter-stage changes, the analysis was extended to compare each stage with its preceding stage. (**A**) EMCI(CN) showed 21 and 15 number of brain regions significant brain regions for male and females, respectively, which was dominated by T1U in males and T2U in females. (**B**) MCI(EMCI) brain regions demonstrated increased dispersion for both perfusion and metabolism in both sexes. (**C**) LMCI(MCI) showed a pronounced increase in cerebral perfusion; however, there was no significant alteration in metabolism over this same interval. (**D**) AD(LMCI) displays several brain regions with characteristics consistent with the NVMF phenotype, where this shift in NVC function is likely a consequence of the neurodegenerative process. (**E-H**) Show the significant regions for each comparison shown in (**A-D**). Brain regions names and indices can be found in Table S4.

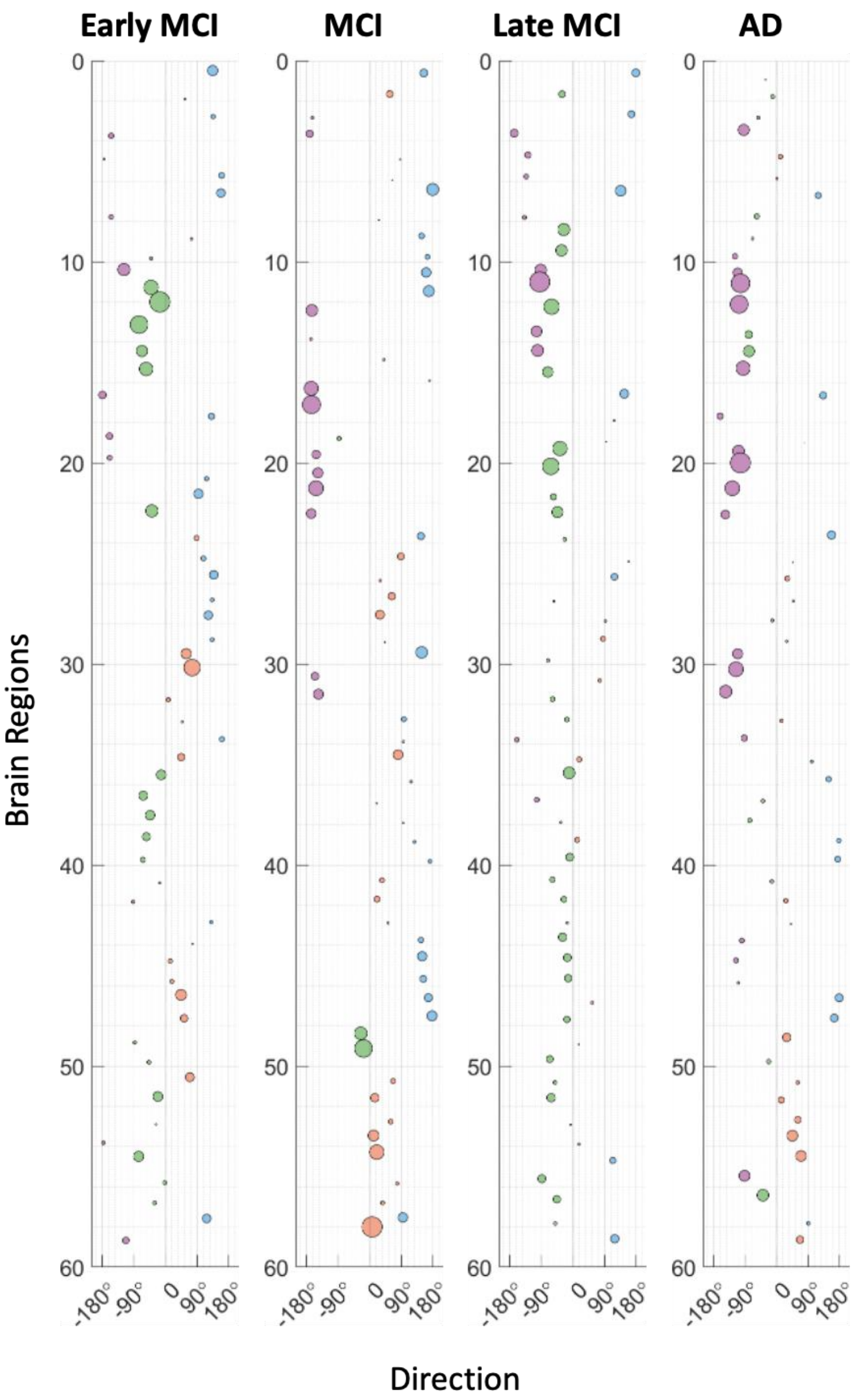


**Supplemental Figure S3 – Complete Continuous Polar Angle Interval Uncoupling Migration Chart Plot for Males Relative to CN.**

In this plot, the origin (0°) corresponds to the reference group; positive movements indicate higher metabolic stages, and negative movements correspond to lower metabolic stages. In this uncoupling migration charts, regions can be seen to be "jumping" between stages, which makes it harder to see the disease progression. Brain regions names and indices can be found in Table S4.

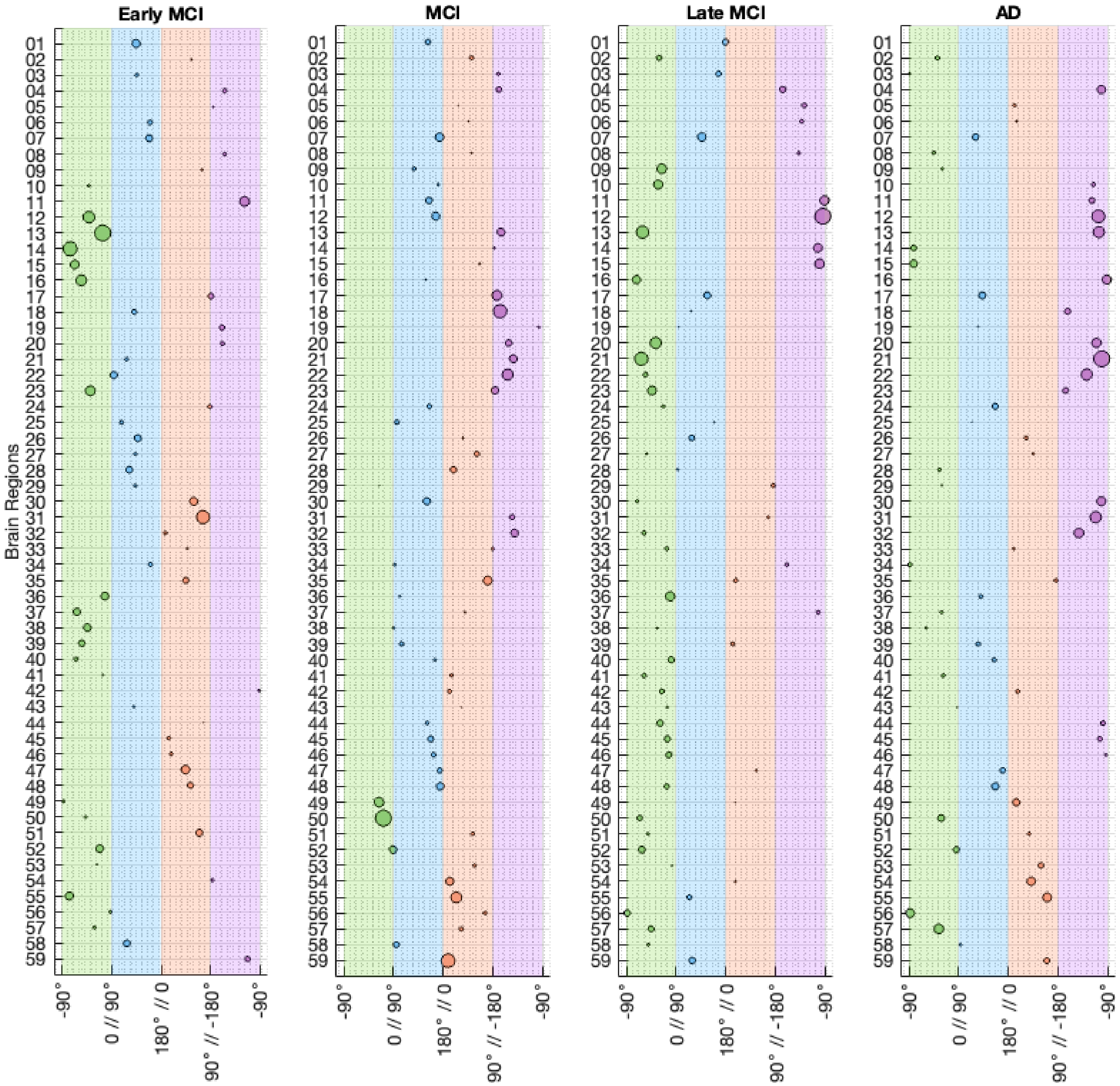


**Supplemental Figure S4 – Complete Uncoupling Migration Chart Plot for Males Relative to CN.**
By sorting the groups by MVD pattern progression, it is easier to see how brain regions progress from one MVD stage to the next one. In addition, it highlights that each brain region progresses at its own rate. Brain regions names and indices can be found in Table S4.

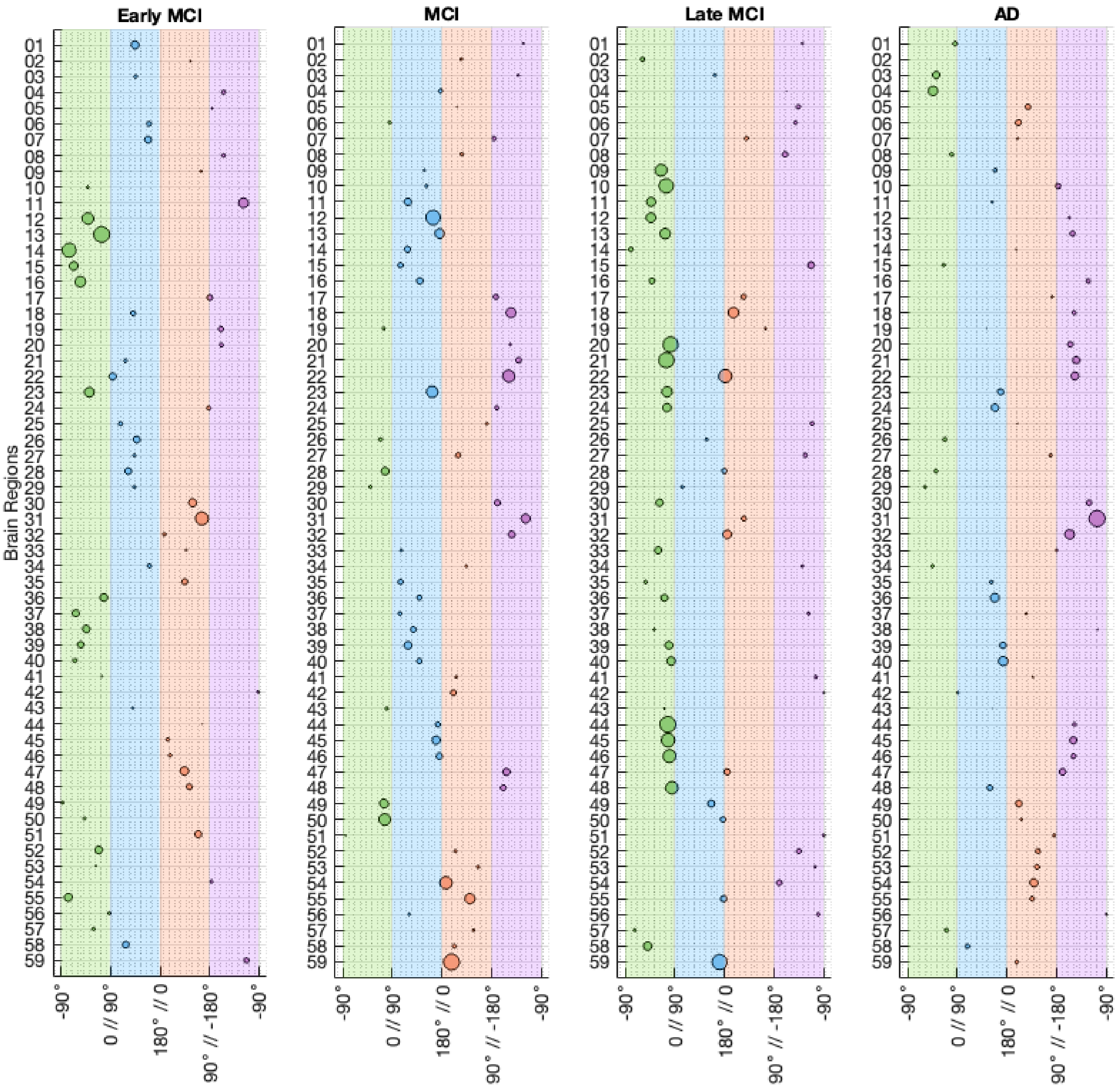


**Supplemental Figure S5 – Complete Uncoupling Migration Chart Plot for Males Relative to the Previous Stage.** Performing the uncoupling migration chart analysis relative to the previous stage highlights the regions progress at different rates. Brain regions names and indices can be found in Table S4.

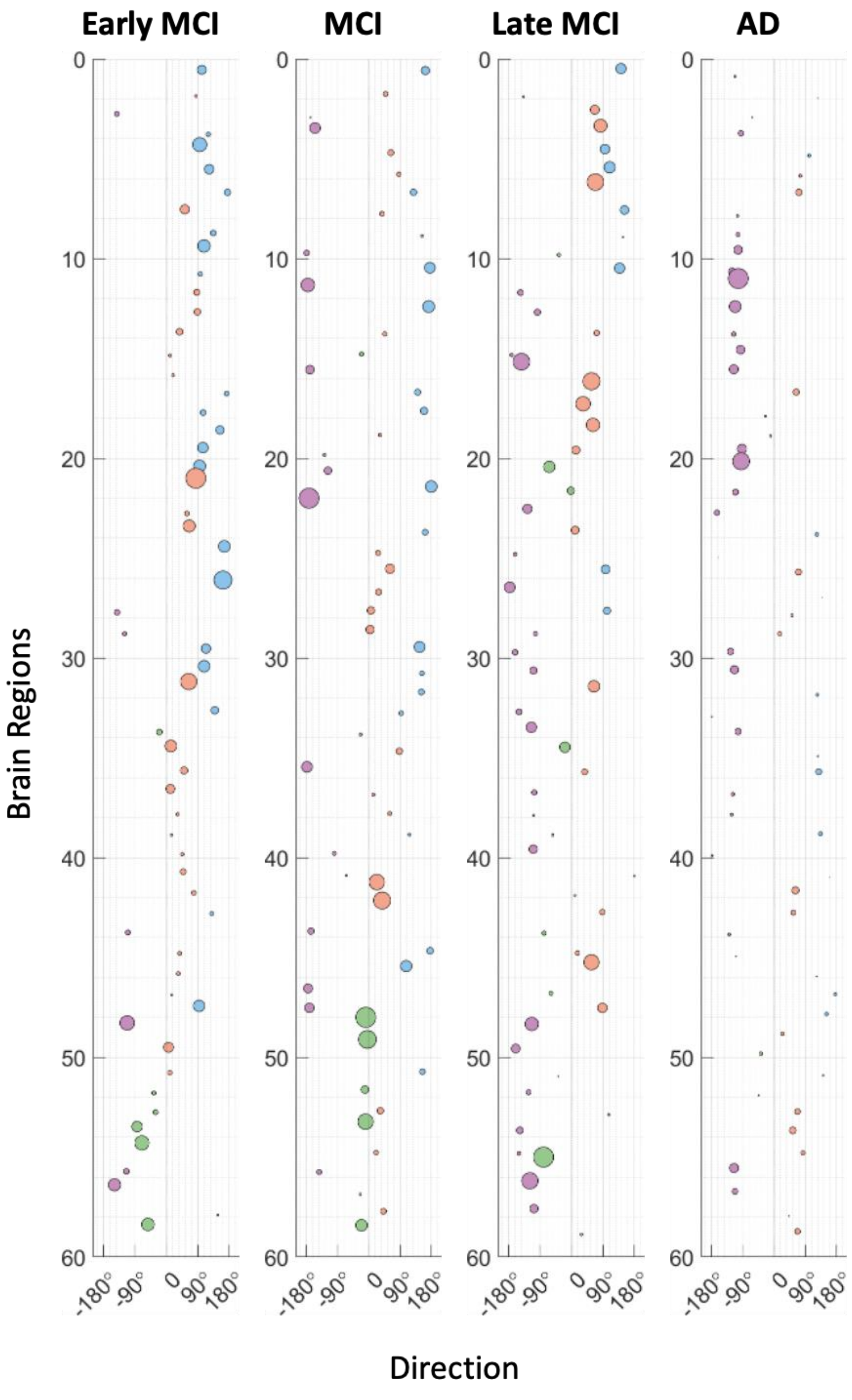


**Supplemental Figure S6 – Complete Continuous Polar Angle Interval Uncoupling Migration Chart Plot for Females Relative to CN.**

The same analysis was performed for female brain regions. The results obtained showed that females progress through the stages at a faster rate. Brain regions names and indices can be found in Table S4.

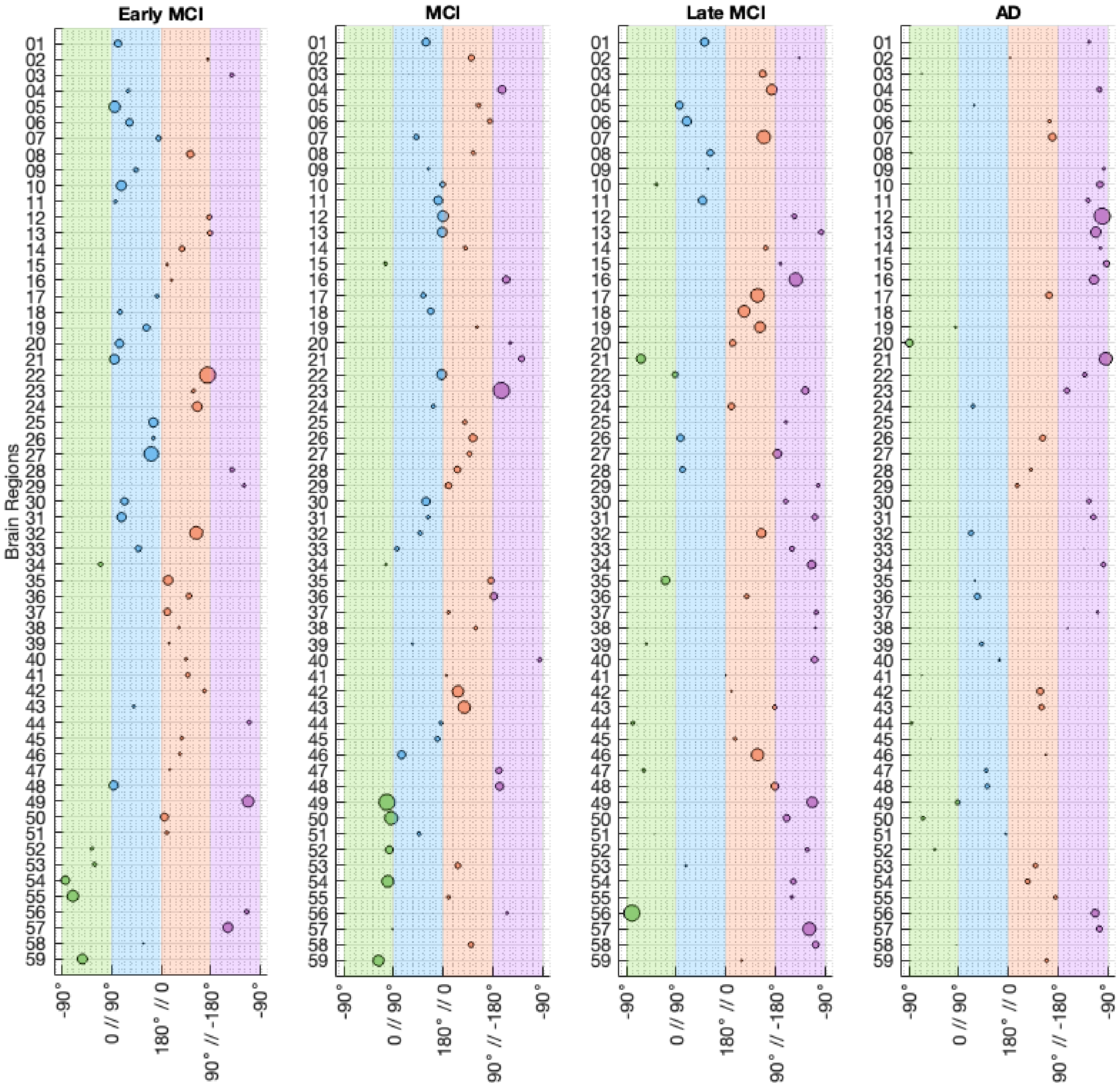


**Supplemental Figure S7 – Complete Uncoupling Migration Chart Plot for Females Relative to CN.**
Similarly to the male population, the same occurs in females when the uncoupling migration charts are arranged by MVD pattern stage. The highlight is that a larger number of brain regions in females already exhibit a T2U phenotype. Brain regions names and indices can be found in Table S4.

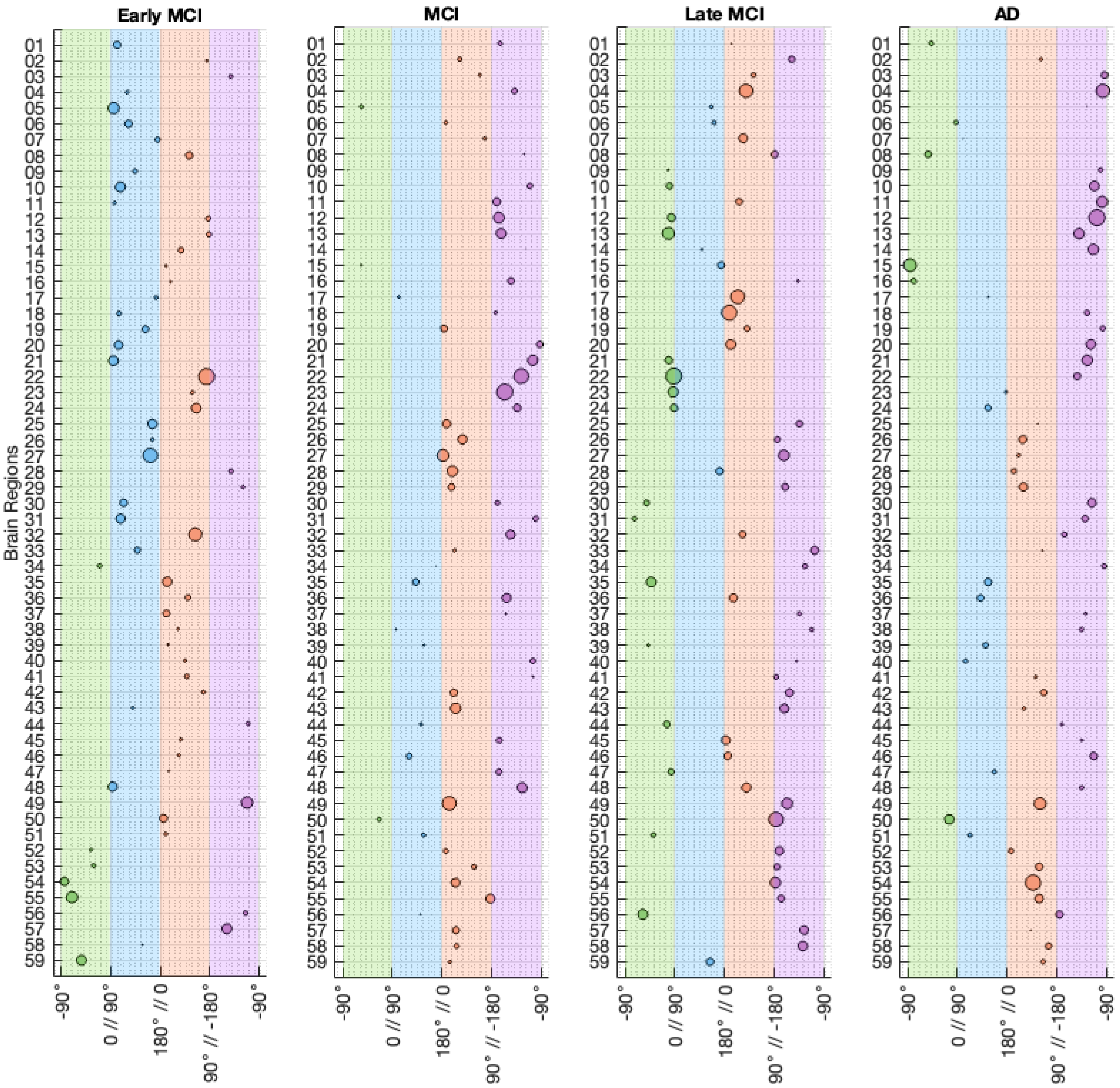


**Supplemental Figure S8 – Complete Uncoupling Migration Chart Plot for Females Relative to the Previous Stage.** Compared to males, analyzing the progression relative to the previous stage in females exhibit the same behavior with the only difference been that female brain regions progress faster than males. Brain regions names and indices can be found in Table S4.

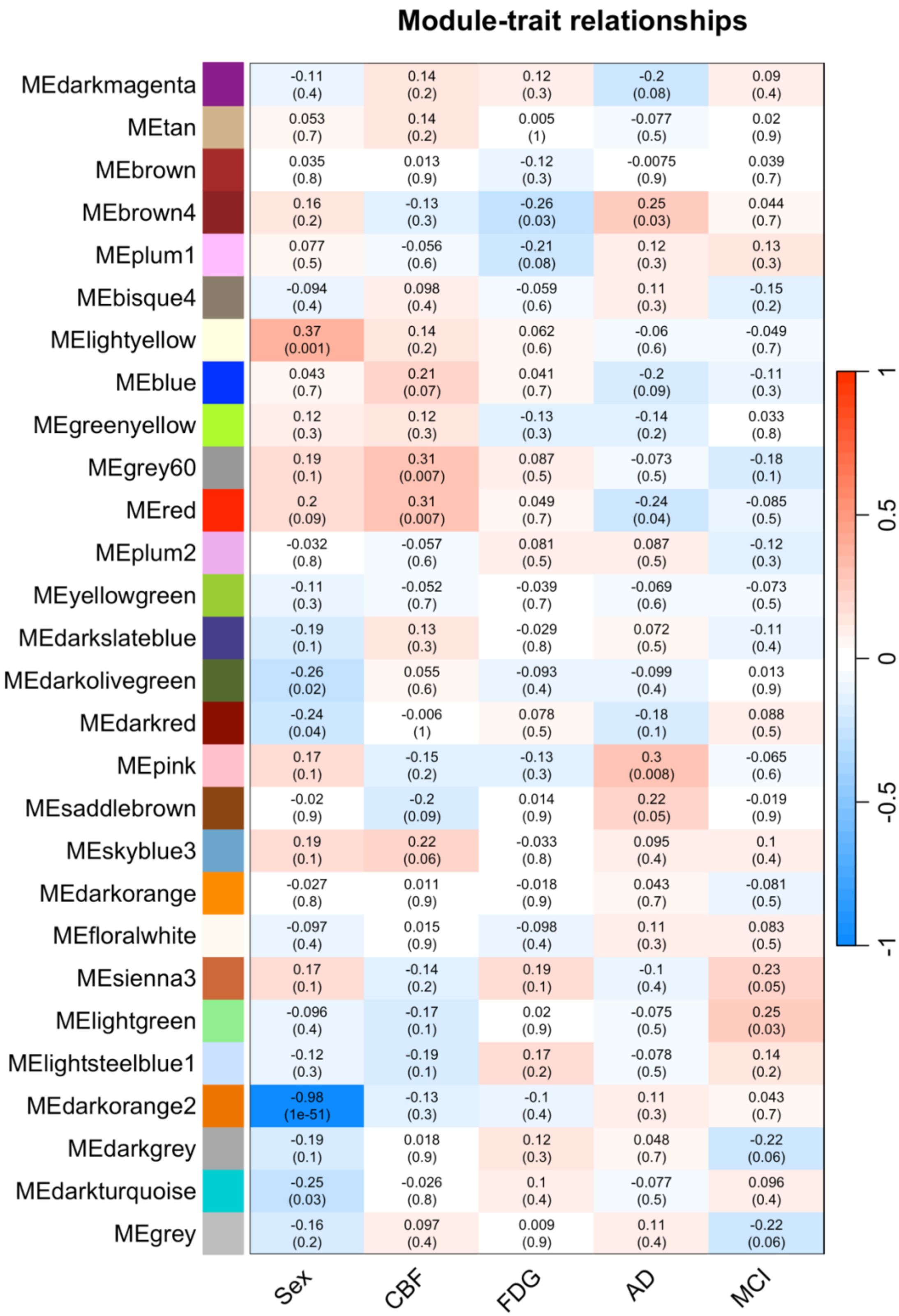


**Supplemental Figure S9 – WGCNA Module Trait Association.**

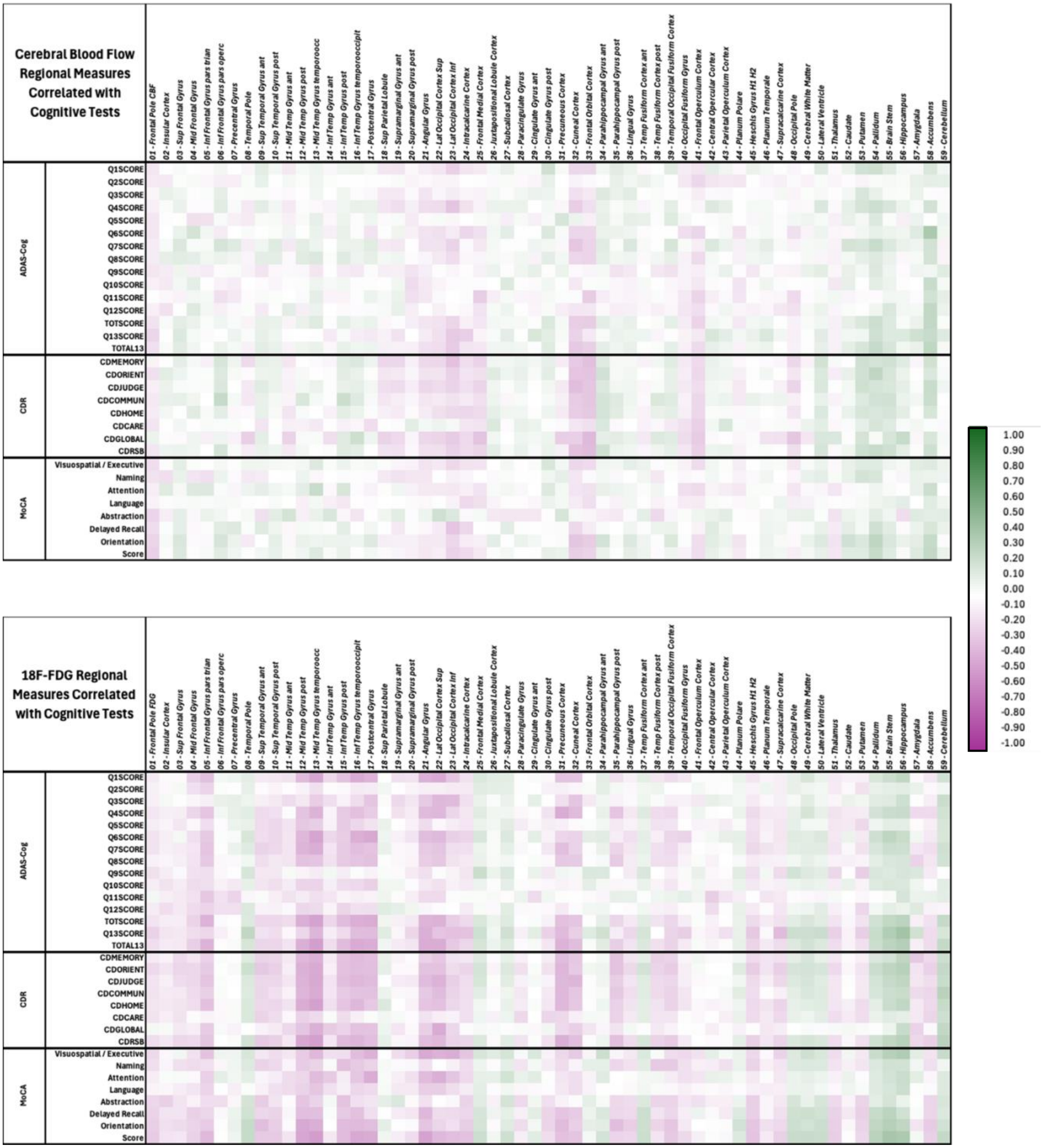


**Supplemental Figure S10 – Correlations of the Clinical Cognitive Assessments with CBF and FDG Measurements for Males.**

By correlating CBF and FDG brain regional measurements with CCA, we aimed to explore the relationship between these variables. For CBF SUVRs, the correlation values for males ranged from -0.33 to 0.36, with a mean of 0.0082±0.0998. This suggests a limited connection between CBF changes and CCA scores. Similarly, for FDG SUVRs, the correlation values ranged from -0.5 to 0.4, with a mean of -0.0551±0.1528. Student's t-tests ($p<0.05$) were employed to determine significant relationships, where most relationships were statistically significant. While the larger intervals indicate a weaker association overall, these correlations still suggest a weak correlation between the imaging measures and CCA. Notably, regions exhibiting positive or negative correlations were consistent across all test sections. These findings imply that CCA may not be sufficiently accurate in capturing the metabolic and perfusion changes associated with disease progression.

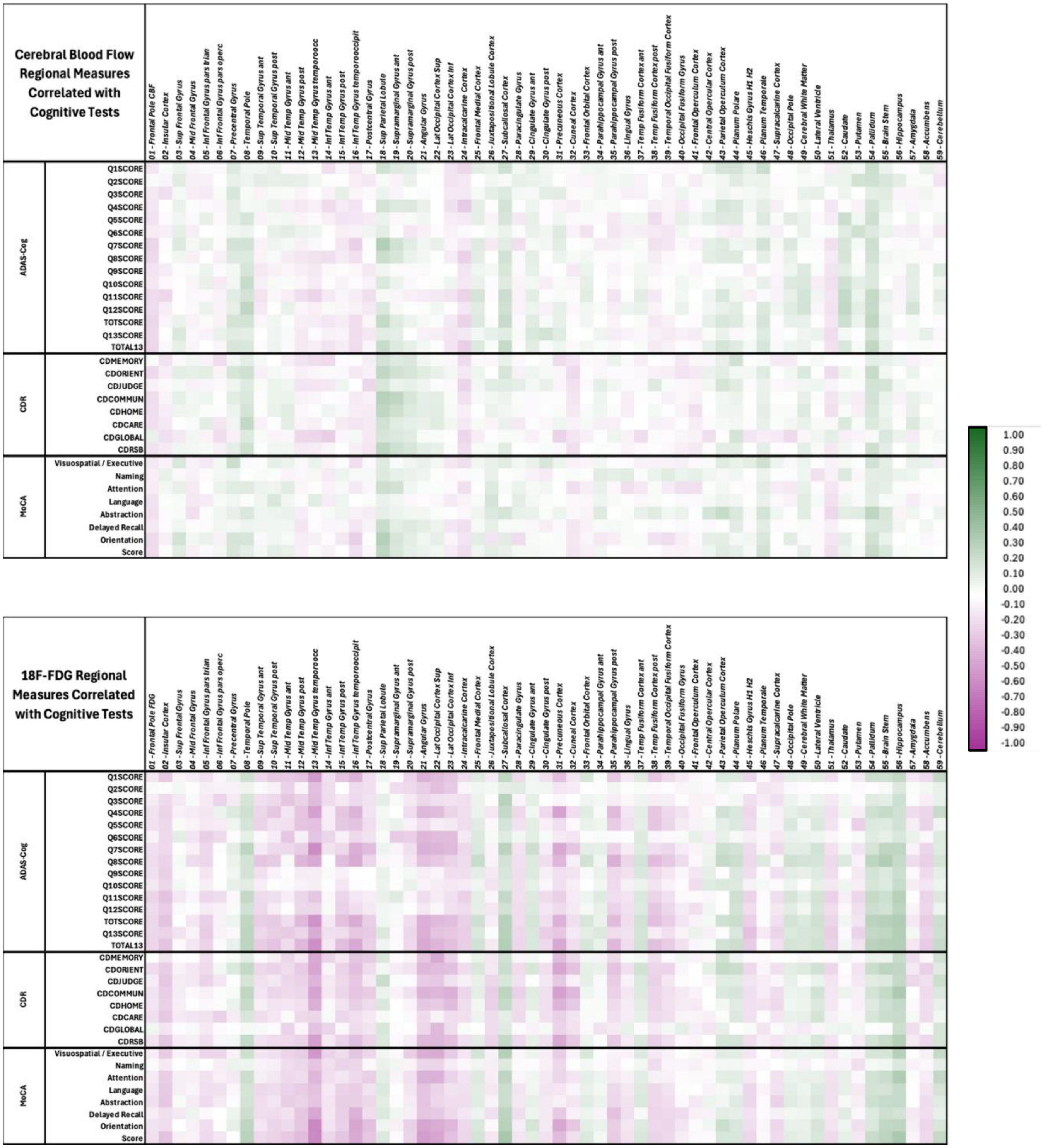


**Supplemental Figure S11 – Correlations of the Clinical Cognitive Assessments with CBF and FDG Measurements for Females.**

Similarly, for females, the correlation values for CBF SUVRs ranged from [-0.27, 0.36], with a mean of 0.0201±0.0917, and the values for FDG SUVRs for females ranged from [-0.6, 0.4], with a mean of -0.0484 ± 0.1594. Using Student's t-tests ($p<0.05$) to determine statistical significance, as in the male case, most relationships were significant. Therefore, these results suggest that the observed behavior in males still apply to females, suggesting that CCA may not be sufficiently accurate in capturing metabolic and perfusion changes associated with disease progression.
